# Hidden pathway for cytokine receptor activation: Structural insights into a marine sponge-derived lectin that activates the thrombopoietin receptor via recognition of the fucose moiety

**DOI:** 10.1101/2021.10.29.466502

**Authors:** Hiromi Watari, Hiromu Kageyama, Nami Masubuchi, Hiroya Nakajima, Kako Onodera, Pamela J. Focia, Takumi Oshiro, Takashi Matsui, Yoshio Kodera, Tomohisa Ogawa, Takeshi Yokoyama, Makoto Hirayama, Kanji Hori, Douglas M. Freymann, Norio Komatsu, Marito Araki, Yoshikazu Tanaka, Ryuichi Sakai

## Abstract

N-glycan-mediated activation of the thrombopoietin receptor (MPL) under pathological conditions has been implicated in myeloproliferative neoplasms induced by mutant calreticulin, which forms an endogenous receptor-agonist complex that constitutively activates the receptor. However, the molecular basis for this mechanism has not been studied because no external agonists existed. We describe the structure and function of a marine sponge-derived MPL agonist, thrombocorticin (ThC), a homodimerized lectin with calcium-dependent fucose-binding properties. ThC-induced activation persists due to limited receptor internalization. The strong synergy between ThC and thrombopoietin suggests that ThC catalyzes the formation of receptor dimers on the cell surface. MPL is subject to sugar-mediated activation, where the kinetics differ from those of cytokines. This result suggests the presence of diverse receptor activation pathways in human thrombopoiesis.

**One-sentence summary:** A marine sponge lectin catalyzes thrombopoietin receptor dimerization and activation, exhibiting strong synergy with thrombopoietin, and modulates internalization of the receptor.

---

The thrombopoietin (TPO) receptor MPL plays critical roles in hematopoietic stem cell (HSC) maintenance and platelet production (*1-3*). MPL, which lacks kinase activity, is activated via Janus kinase 2 (JAK2) bound to the intracellular domain of MPL. The detailed process of receptor dimerization and activation of JAK2 by TPO is poorly defined, partially due to the lack of structural information on MPL (*4*). Under pathological conditions, myeloproliferative neoplasms (MPNs), a mutant form of the glycan-dependent molecular chaperone CALR (CALRmut), bind to the immature sugar chain of MPL in the endoplasmic reticulum and then translocate to the cell membrane to form a complex with functional MPL (*5-7*). This complex leads to the transformation of hematopoietic cells in an MPL-dependent manner (*8-10*). CALRmut activates MPL only when CALRmut binds to the immature sugar chain of the receptor (*10, 11*). However, the external agonist required for the recapitulation of this mode of activation is not known. Therefore, the molecular basis of receptor activation by lectin-type ligands remains largely unstudied.

We recently identified a marine sponge-derived 14-kDa protein, thrombocorticin (ThC), as a potent agonist of MPL (*12*). We report the three-dimensional structure of ThC as a novel fucose-binding lectin and the mechanisms underlying its MPL activation by binding to sugar chains on MPL.

## Biochemical profiles of ThC

We first identified a preliminary 131-amino acid sequence of native ThC (nThC) isolated from the sponge using mass spectrometry and Edman degradation after peptic digestion (Fig. 1A (i) and Fig. S1). A draft ThC based on this sequence assigned lysine (K) to the 25^th^ amino acid residue. However, unlike nThC (*12*), the draft ThC failed to promote the proliferation of MPL activation-dependent Ba/F3-HuMpl, even at high concentrations (Fig. 1B). This lack of activity showed that at least one amino acid residue in the draft-ThC sequence was assigned incorrectly. Therefore, we performed X-ray crystal structure analysis of nThC and found that the 25^th^ amino acid residue originally assigned as K in the draft-ThC sequence was glutamic acid (E) or glutamine (Q) (Fig. S2). An LC-MS/MS experiment unambiguously identified this residue as Q (Fig. S3), and the complete amino acid sequence of ThC was determined.

**Fig. 1.**
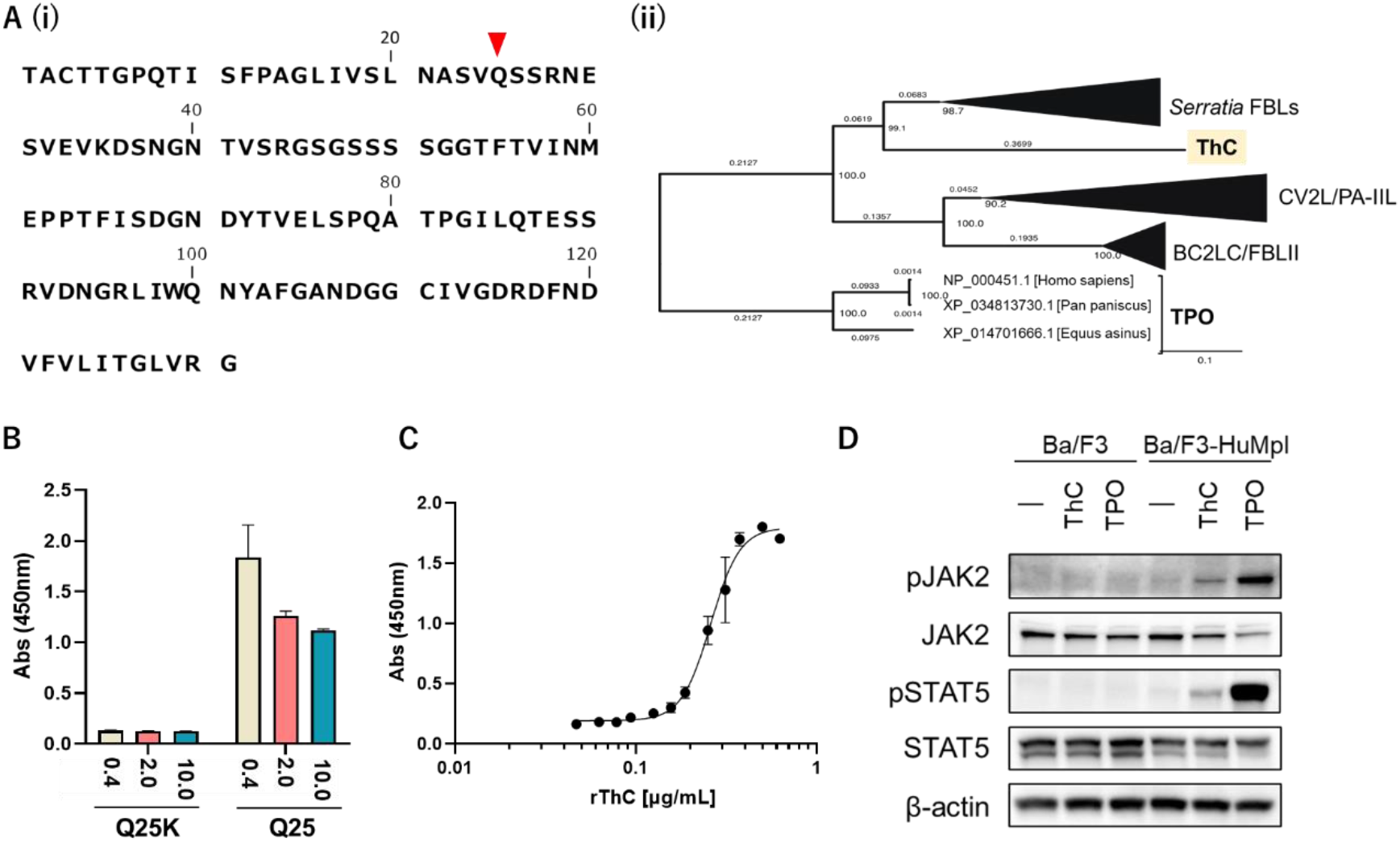
Biochemical profiles of ThC. (A) (i) Amino acid sequences of ThC. The red arrowhead indicates the Q residue at position 25, which was assigned as K in the draft sequence. (ii) Phylogenetic tree of ThC, related bacterial lectins and TPO with collapsed tree nodes (Fig. S5). (B) A comparison of the activity of draft ThC (Q25K) and rThC (Q25) in Ba/F3-HuMpl cell proliferation. Values shown at the bottom indicate concentration of added rThC and draft ThC (μg/mL). (C) Concentration-response curve of Ba/F3-HuMpl cells proliferating by recombinant ThC (rThC). The half-maximal effective concentration EC_50_ was 0.26 and 0.31 μg/mL for rThC and nThC (*12*), respectively. (D) Immunoblot analysis of Ba/F3 and Ba/F3-HuMpl cells upon steady-state activation by TPO and rThC.

*N*-terminal His-tagged recombinant ThC (rThC) harboring Q at position 25 promoted the proliferation of Ba/F3-HuMpl cells (Fig. 1C) in an MPL-dependent manner (Fig. S4). Immunoblot analyses indicated that rThC activated steady-state JAK/signal transducer and activator of transcription (STAT) signaling (Figs. 1D). These data indicated that rThC activated MPL to promote cell proliferation in Ba/F3-HuMpl cells.

## Critical role of sugar-binding capacity in ThC-dependent MPL activation

The amino acid sequence of ThC shares approximately 33% identity with bacterial fucose-binding lectins. However, little similarity was found between ThC and TPO in amino acid sequences (Fig. 1A (ii)) and three-dimensional structures. The activation of MPL by lectin-like molecules that are structurally distinct from TPO attracted our attention, because the CALRmut interaction with an immature sugar chain attached to the receptor during receptor maturation was proposed as an alternative mode of pathological MPL activation in MPN (*5, 7*). Therefore, we hypothesized that *ThC binds to sugar chains on the extracellular domain of the receptor on the cell surface to promote MPL activation*. Consistent with this hypothesis, L-fucose or D-mannose inhibited ThC-dependent cell proliferation in Ba/F3-HuMpl cells (Fig. 2A). In contrast, the effects of the sugars on TPO-dependent cell proliferation were negligible (Fig. S6). The effects of fucose and mannose were concentration-dependent, with half-maximal inhibitory concentration (IC_50_) values of 22.7 and 6,667 μM, respectively (Fig. 2B). Subsequent sugar affinity column experiments showed that ThC bound to fucose and mannose only in the presence of Ca^2+^ (Fig. 2C). Isothermal titration calorimetry (ITC) experiments confirmed Ca^2+^ dependence, with binding dissociation constants (K_D_s) of rThC bound to fucose or mannose in the presence of 5 mM Ca^2+^ of 4.72 and 66.2 × 10^−6^ M, respectively, which is consistent with the observations described above (Fig. 2D and Table S1). Taken together, these data suggested that blockade of ThC binding to MPL sugars inhibits ThC-dependent cell proliferation.

**Fig. 2.**
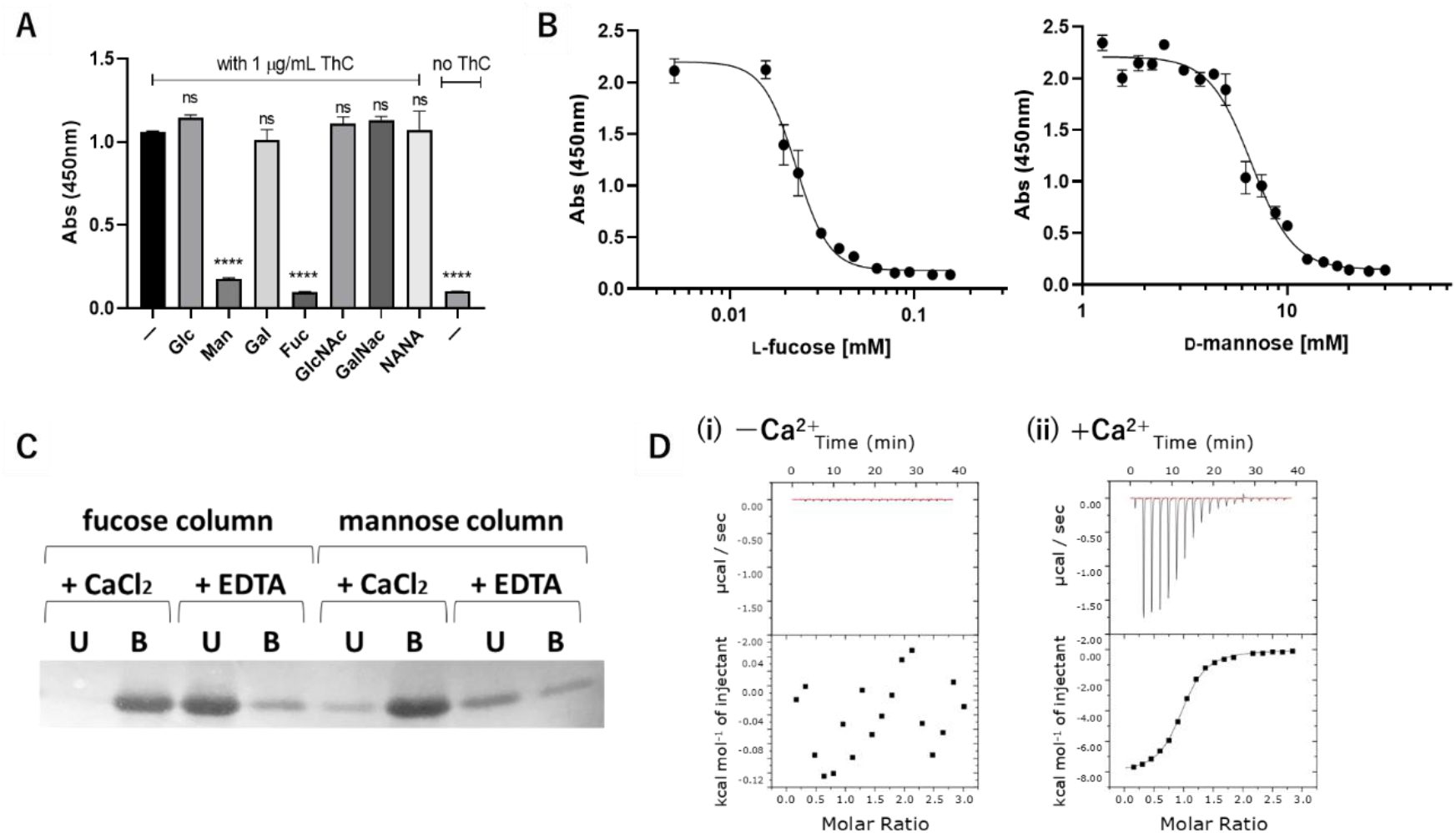
Critical role of sugar-binding capacity in ThC-dependent MPL activation. (A) Relative cell proliferation in the presence of various sugars (10 mM) in Ba/F3-HuMpl cells with rThC (1 μg/mL). \*\*\*\**P* < 0.001, ns, not significant. (B) Concentration-dependent inhibition of fucose or mannose on cell proliferation by rThC treatment (1 μg/mL). (C) Binding capacity of rThC to fucose- or mannose-immobilizing resins in the presence of 1 mM CaCl_2_ or 1 mM EDTA. Unbound (U) and bound (B) rThC to the resin is shown as a pair. (D) Thermodynamic analysis of the interaction with fucose in the absence (left) and presence (right) of 5 mM CaCl_2_ using ITC. Thermogram (top) and titration curve (bottom) are shown.

## Structural insights of ThC as a homodimeric lectin

To aid in the determination of the primary structure of ThC and gain further structural insights into ThC itself, we determined the crystal structure of nThC at 1.6 Å resolution (Fig. 3). ThC has a *β*-sandwich structure composed of nine *β*-strands. An intramolecular disulfide bridge is formed between Cys3 and Cys111 (Fig. 3A), and two ThC molecules assemble as a homodimer in the crystal (Fig. S7 and 3B). This structural feature is shared with other fucose-binding lectins, *i*.*e*., C-terminal domain of BC2L-C (BC2L-C - CTD) and PA-IIL, which are homologous proteins of ThC, and suggests that dimeric assembly is an important shared structural characteristic (Fig. S7). The structure of rThC superposed well onto nThC (root-mean-square deviation (r.m.s.d.) of 0.66 Å for 261 Cα atoms), which suggested that it was structurally and functionally equivalent to nThC. Therefore, His-tagged rThC was used for further structural, physico-chemical, and physiological analyses, and it is termed ThC hereafter (Fig. S8). Structural insights into ThC, particularly the formation of homodimers, provided a rational model for MPL activation that was likely triggered by the homodimerization of receptor molecules.

**Fig. 3.**
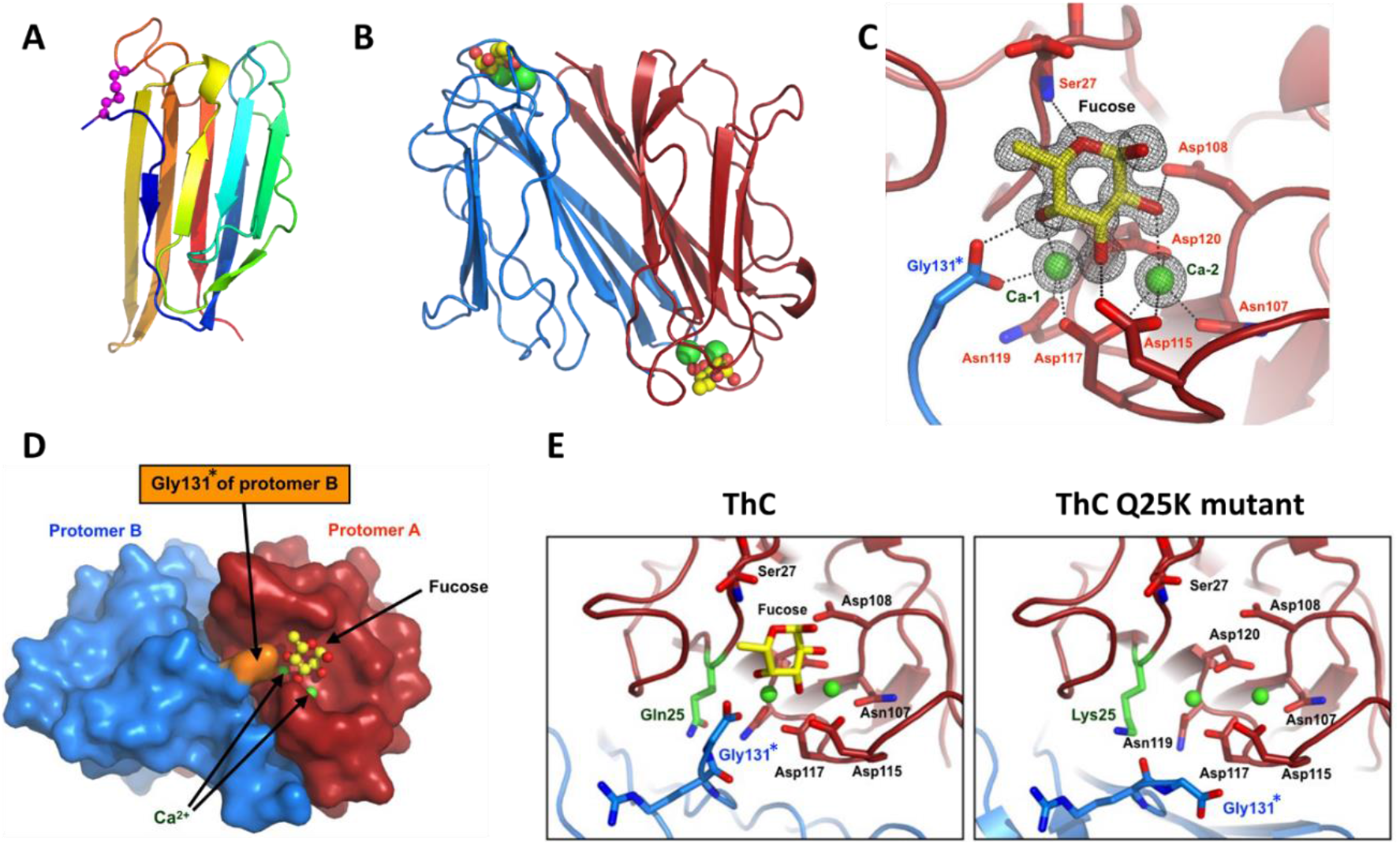
Crystal structure of ThC. (A) Ribbon diagram of the nThC monomer colored according to the sequence in blue at the N-terminus to red at the C-terminus. The disulfide bond between Cys3 and Cys111 is shown as a magenta ball. (B) Dimer structure of rThC in complex with Ca^2+^ (green ball) and fucose (ball-and-stick model, yellow: carbon, red: oxygen). (C) A close-up view of the fucose binding site of rThC. The bound Ca^2+^ ions and fucose are shown as green balls and stick models, respectively. Residues are colored according to the chain as in (B). A Fo-Fc map of the fucose and Ca^2+^ is shown. (D) Pseudodomain swapping structure in dimeric rThC. Gly 131*, shown in orange, of one protomer intervenes in the other. (E) Structural comparison of the Ca^2+^-binding configuration between wild type (left) and Q25K (right). Close-up view of the fucose binding site is shown. Individual protomers are shown in red and blue. The substituted residues (Q25 and K25) are shown in green.

## Structural basis for ThC sugar binding

The crystal structure of ThC in complex with L-fucose (Fig. 3C) or D-mannose (Fig. S9) showed that the sugar molecule was bound in a cavity of the dimeric protein at the interface between two protomers via two Ca^2+^ ions, Ca-1 and Ca-2. Ca-1 was chelated in a polar cavity formed by D117-N119-D120 of one protomer and the C-terminal carboxylate of G131 of the other protomer (denoted as Gly131*, Fig. 3C). This unique structural feature, herein called the ‘pseudodomain-swapping motif’, was formed between the protomers, and it is a characteristic hallmark of this protein family (Fig. 3D). Ca-2 was chelated by polar groups along the N107-D115-D117-D120 sequence (Fig. 3C). Sugar is recognized by ThC via polar interactions along the Ca-1-Ca-2-D115-D120-D108 sequence of one protomer and three hydroxy groups of the carbohydrates. Notably, the pseudodomain-swapping structure enables stable carbohydrate binding via a hydrogen bond network established between the carboxylate of G131* of the adjacent protomer and Ca-1 and O4 of fucose (O2 of mannose). The positions of the carbohydrates were further stabilized by binding between O5 and Ser27 in both sugars. This manner of recognition shows the importance of Ca^2+^ in carbohydrate binding, which is consistent with the Ca^2+^ dependency of carbohydrate binding of ThC (Fig. 2C, D and Table S1). Stereochemical orientation of the three hydroxyl groups is shared between L-fucose and D-mannose but not the other chain-bearing sugars tested, which explains the carbohydrate specificity of ThC (Fig. S10). A 1:1 stoichiometry between ThC and mannose/fucose was apparent in the ITC analysis (Fig. S11).

In the draft sequencing study of ThC, we coincidentally found that Q25 was a key residue for its agonist action (Fig. 1B). The ITC data showed a complete lack of affinity of Q25K (draft-ThC) for fucose (Fig. S11). To examine its structural basis, the Q25K mutant was crystallized in the presence of Ca^2+^. The C-terminus in draft-ThC faced away from Ca-1. The structure of draft-ThC clearly differed from nThC in the conformation of the C-terminus in the counterpart protomer and the position of the side chain of Q25 (Fig. 3E). This conformational change caused the loss of the pseudointerprotomer swapping domain and resulted in loss of Ca-1 coordination of the C-terminal carboxylate group of G131*, leading to the profound loss of fucose-binding capability and agonist activity of the mutant (Fig. 1B). We confirmed this series of changes by preparing a G132 mutant in which an extra G residue was added to the C-terminus to alter the pseudodomain-swapping motif. As expected, ITC analysis revealed that the G132 mutation exhibited diminished fucose-binding activity (Table S1 and Fig. S11), and agonist activity was completely lost (Fig. S12). These observations led to the identification of the structural determinants for the sugar-binding and agonist actions of ThC. Specifically, the binding cavity of one protomer, two calcium ions, and the C-terminal domain of the other protomer together stabilize the sugar-bound state of the protein.

## Activation of MPL by ThC via a fucosylated sugar chain

To gain further insight into the sugar-mediated activation of MPL, we assessed the effects of lectins bearing fucose- or mannose-binding properties on the proliferation of Ba/F3-HuMpl cells (Fig. 4A). None of the lectins tested, except PA-IIL (*13, 14*), a fucose-binding lectin homologous to ThC, promoted the proliferation of Ba/F3-HuMpl cells. Despite the high degree of structural similarity between ThC and PA-IIL (r.m.s.d. 2.05 Å for 104 Cα atoms, Fig. S7), PA-IIL showed approximately 70-fold reduced potency in inducing MPL-dependent cell proliferation compared with ThC (Fig. 1C and 4B).

**Fig. 4.**
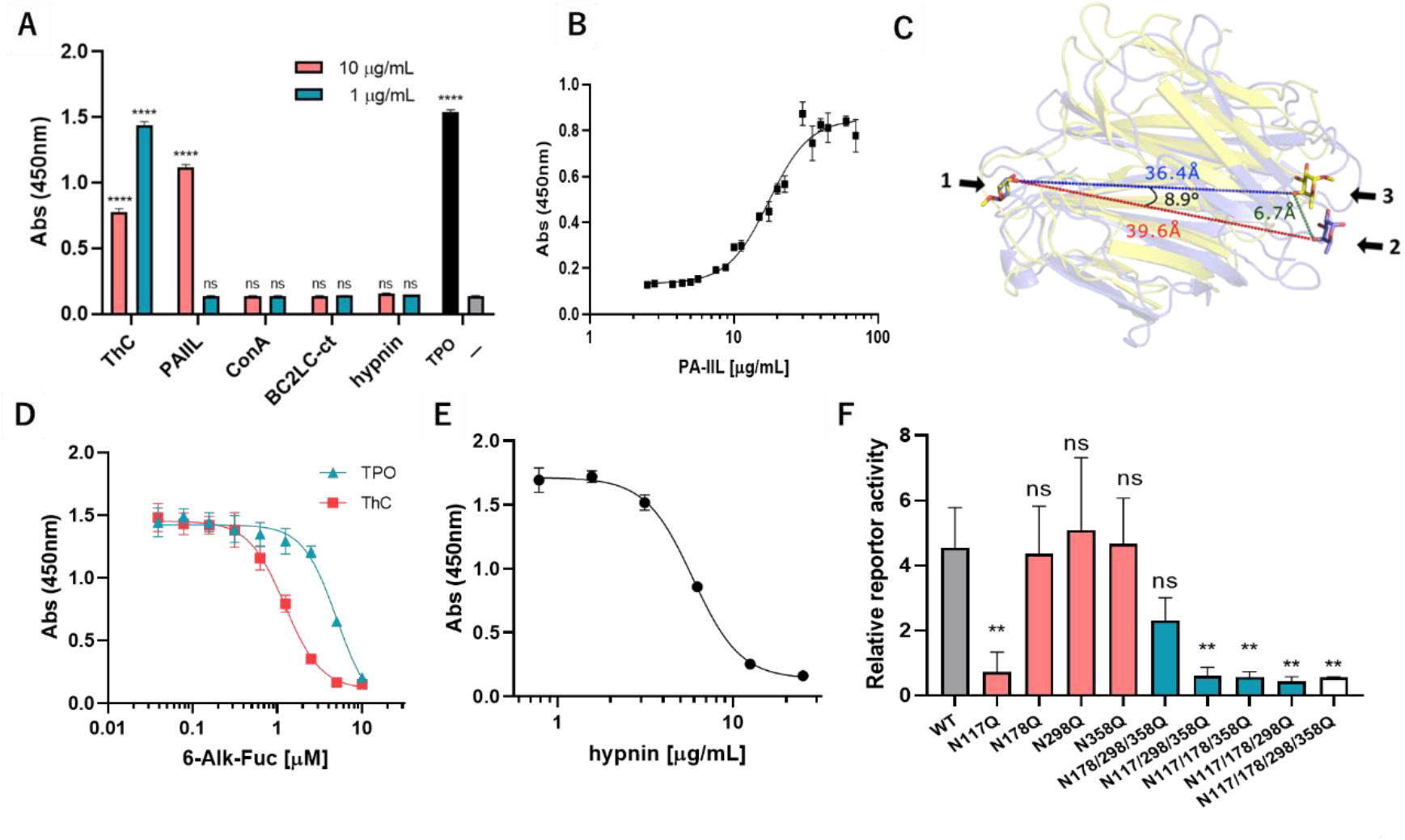
Involvement of the fucose moiety in ThC-dependent MPL activation. (A) Effect of lectins on the proliferation of Ba/F3-HuMpl cells: PA-IIL, a fucose/mannose selective bacterial lectin with high homology to ThC; BC2L-C-CTD, mannose-selective bacterial lectin with high homology to ThC; hypnin, a core 1,6-fucosylated glycan-specific algal lectin; ConA, a mannose-specific legume lectin. TPO, 10 ng/mL; (-), no agonist. (B) Concentration dependency of Ba/F3-HuMpl cell proliferation induced by PA-IIL (EC_50_ = 17.5 μg/mL). (C) Relative positions of two fucose molecules bound to the ThC dimer and PA-IIL dimer. One of the two fucose molecules bound to each is superimposed (arrow 1). The other fucose molecule is shown as sticks. Purple (arrow 2) represents the fucose bound to ThC, and yellow (arrow 3) represents the fucose bound to PA-IIL. The numbers represent the distance between O4 atoms (red: between two fucoses of ThC, blue: between two fucoses of PA-IIL, green: between fucoses of ThC and PA-IIL) and the angle between the three O4s of superimposed fucoses, of the PA-IIL-bound fucose, and the ThC-bound fucose. Ribbon diagrams of the ThC dimer (purple) and PA-IIL dimer (yellow) are also shown in translucent form. (D) Preferential inhibition of ThC-dependent cell proliferation in Ba/F3-HuMpl cells by 6-alkynyl-fucose. Cells were cultured in the presence of ThC (1 μg/mL) or TPO (10 ng/mL). (E) Hypnin-mediated inhibition of cell proliferation induced by ThC (1 μg/mL), with an IC_50_ value of 5.88 μg/mL. (F) Effect of potential N-glycosylation site MPL mutations on activation by ThC (1 μg/mL). STAT5 reporter activity representing MPL activation status is depicted. Gray bar: wild-type (WT) MPL; red bars: single-site mutant MPL; blue bars: triple-site mutant MPL; and white bar: quadruple-site mutant MPL. \*\**P* < 0.01, \*\*\*\**P* < 0.001, ns: not significant.

This result suggested that the dimerization and sugar specificity of a lectin alone are insufficient for an MPL agonist, although certain structural features inherent to ThC and PA-IIL contributed to their agonist action. Lack of agonist activity by BC2L-C-CTD, which belongs to the same family as PA-IIL (r.m.s.d. 1.18 and 2.13 Å for 111 and 106 Cα atoms with PA-IIL and ThC, respectively), strongly suggested that sugar-binding properties, specifically fucose binding, are crucial for MPL activation in addition to structural similarity. The ITC data of PA-IIL and BC2L-C-CTD clearly showed that PA-IIL bound to fucose and mannose (*15*), and BC2L-C-CTD bound only to mannose (Fig. S13 and Table S1). These data support the importance of fucose binding activity for MPL activation. Notably, MPL activation by PA-IIL was much weaker than that by ThC (Fig. 4B), although the fucose binding affinity of PA-IIL was stronger than that of ThC (Table S1). Therefore, we determined the inherent structural differences between the lectins by comparing the positions of two fucose molecules bound to ThC and PA-IIL. The distance between the representative atom of fucose O4 was 39.6 Å for ThC and 36.4 Å for PA-IIL (Fig. 4C). When one fucose molecule was superimposed, the position of the other was shifted by approximately 6.7 Å, such that the angle between three O4 atoms was 8.9 º (Fig. 4C).

To demonstrate the importance of the fucosylated sugar chain, we treated Ba/F3-HuMpl cells with peracetylated 6-alkynyl fucose (6-Alk-Fuc), an inhibitor of GDP-fucose synthase (TSTA3), which attenuates the formation of fucose-containing sugar chains (*16*). The resulting 6-Alk-Fuc Ba/F3-HuMpl cells were treated with ThC or TPO. 6-Alk-Fuc decreased ThC- and TPO-induced cell proliferation, with IC_50_ values of 1.2 and 5.0 μM, respectively (Fig. 4D). These results indicated the considerable contribution of fucosylated glycans to receptor activation. However, the type of fucosylated chain interacting with ThC could not be specified via an analysis of the aforementioned data alone. Human FUTs catalyze α(1,2)-, α(1,3)-, α(1,4)-, α(1,6)-, and *O*-fucosylation, and cell surface glycans may exhibit any of these fucosylation patterns (*17*). Therefore, we tested the effect of hypnin, an algal lectin with highly strict recognition of core α(1,6)-fucosylated glycans (*18*), on the action of ThC. We found that hypnin inhibited ThC-induced cell proliferation in a concentration-dependent manner, with an IC_50_ of 5.88 μg/mL (Fig. 4E). Because the cytostatic concentration of hypnin was much higher (approximately 47 μg/mL, Fig. S14), this result was ascribed to competitive inhibition between ThC and hypnin for core α(1,6)-fucose (Fig. 4E).

We assessed the actions of ThC against glycan mutants of MPL. The extracellular domains of MPL have four consensus amino acid sequences for *N*-type glycans at N117, N178, N298, and N358 (*19*)(Fig. 4F). To determine the critical site for ThC activation, mutants in which N residues were replaced by Q residues were expressed in HEK293T cells, and receptor activation was monitored using the STAT5 reporter assay. The mutations did not significantly affect receptor activation by TPO (Fig. S15), showing that the mutation itself had little effect on the receptor activation ability. However, differential activation between the mutants was apparent with ThC application (Fig. 4F). Notably, the N117Q mutant significantly lost sensitivity. Three of the four consensus N residues were replaced with Q residues, which left only one glycan. These mutations significantly suppressed ThC activation, but the mutant with N178/298/358Q, in which the N117 sugar remained, responded to the treatment. These data clearly support the supposition that the TPO receptor may be activated via ligand binding at the surface glycan attached to the N117 residue of MPL.

Notably, the glycan site identified here is the same site that was previously attributed to activation by CALRmut under pathological conditions (*5-7, 20*). Because the property of CALRmut for MPL remains largely elusive due to a lack of an assay system (see above), we examined the mode of ThC-induced receptor activation. Unlike TPO, which induced the activation of MPL in 10 min and resulted in a rapid attenuation of activation, ThC activated MPL at 30 min after ligand addition and exhibited a capacity for prolonged activation (Fig. 5A). When the levels of accumulated cell surface receptors were measured upon activation with a set of agonists, the receptors remained on the cell surface in the ThC-treated cells (Fig. 5B). This differed from the receptors in TPO-treated cells, in that the receptor population gradually decreased due to internalization. Likewise, prolonged accumulation of MPL on the cell surface was observed in CALRmut-expressing cells, where receptor activation was persistent (*5*). These observations suggest that the mode of activation by lectin-type ligands is slow and steady, but cytokine-mediated activation is rapid and extinctive.

**Fig. 5.**
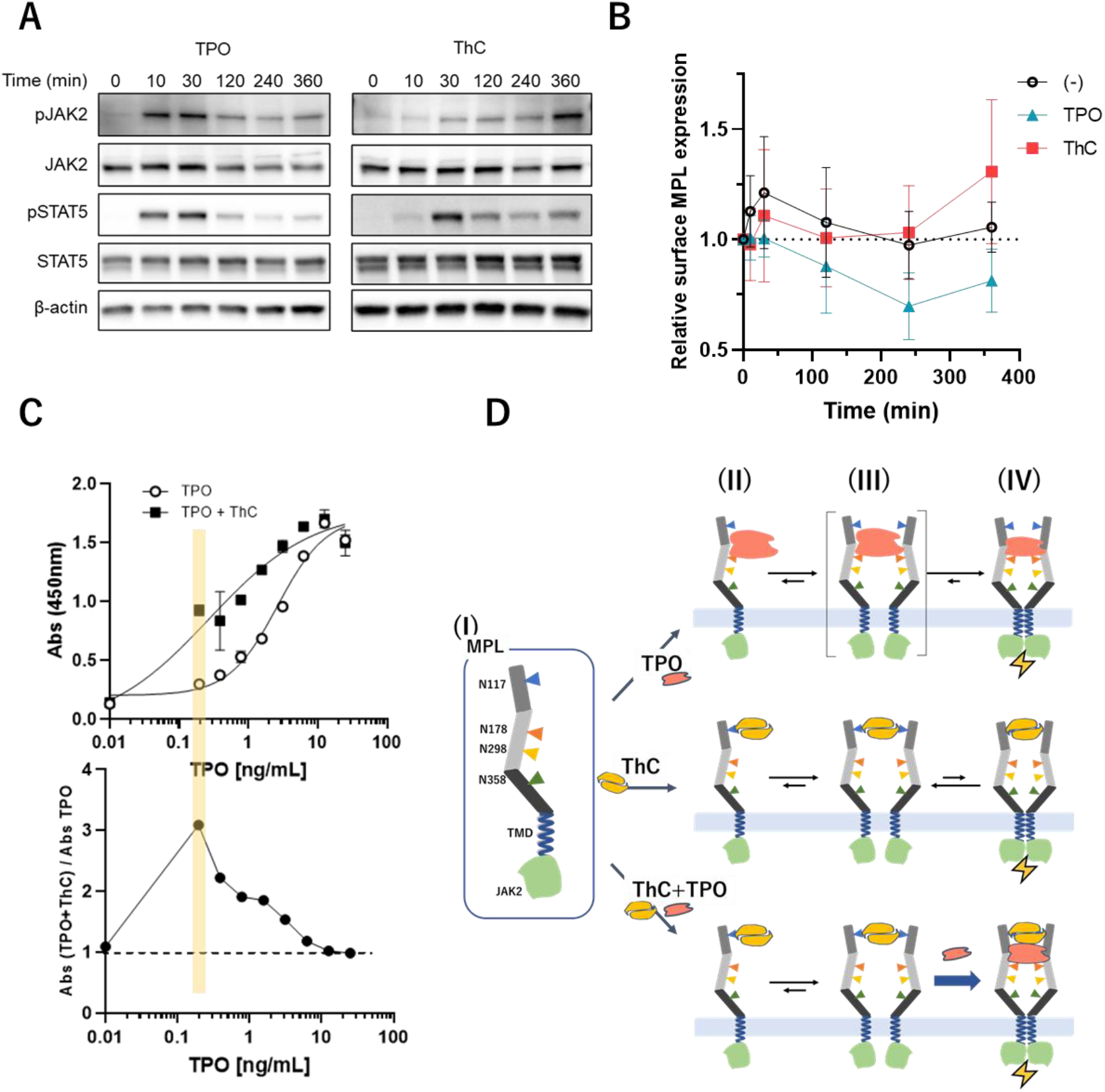
Differential processes of MPL activation. (A) Time-dependent activation of MPL-downstream molecules in Ba/F3-HuMpl cells treated with TPO (3 ng/mL) and ThC (1 μg/mL). Phosphorylation of JAK2 and STAT5 was monitored at the indicated times after the addition of an agonist. (B) Relative amount of MPL on the cell surface 0, 10, 30, 120, 240, and 360 min after the addition of ThC (1 μg/mL) and TPO (3 ng/mL). (C) Synergistic effects observed in each combination of agonists in the Ba/F3-HuMpl cell proliferation assay. Concentration-response curves for tested agonists in the absence or presence of fixed subactivation concentrations of ThC (0.1 μg/mL) (upper trace). The ratios of absorptions with and without ThC (0.1 μg/mL) are plotted (lower trace). (D) Proposed mechanisms of activation by two mechanistically discrete agonists, TPO and ThC: (I) Schematic depiction of MPL. Monomeric MPL has four N-glycosylation sites at N117, 178, 298, and 358; (II) ligand-bound monomeric state with each TPO and ThC; (III) ligand-bound nonactivated state; (IV) dimeric signaling complexes.

## Potentiation of TPO activity by lectins

Our observations strongly support the hypothesis that *MPL can be activated by a completely different mechanism from currently known agonists*. Because lectin-type ligands likely maintain increased levels of receptors on the cell surface, we assessed whether these ligands synergized with natural ligands. We examined the concentration-dependent proliferation of TPO in Ba/F3-HuMpl cells in the presence of a subactivating concentration of ThC (0.1 μg/mL). As expected, ThC synergistically enhanced the agonist action of TPO (Fig. 5C). The rate of enhancement had a bell-shaped relationship with TPO concentration and showed the greatest effect when 0.2 ng/mL TPO was applied (Fig. 5C, lower tracing). At this concentration, TPO alone induced only a 10% increase in cell proliferation but became 50% in the presence of ThC. The EC_50_ value (ng/mL) of TPO/ThC (0.27) was 10 times lower than that of TPO (2.6). These data supported the idea that bidentate glycan-binding ligands allosterically sensitized the action of TPO.

## Discussion

We determined the three-dimensional structure of the potent MPL agonist ThC as a homodimeric complex of lectin molecules featuring two calcium ions that form a cage-like complex for selective binding to fucose and mannose (Fig. 3C). We showed that ThC bound to fucose and mannose (Table S1) and demonstrated that the fucose-binding property was critical for MPL activation (Fig. 4A, D, E). The mode of MPL activation by ThC resembled the pathogenic ligand CALRmut because of the dependency of the N-glycosylation site on activation (Fig. 4F) and the persistent accumulation of receptor molecules on the cell surface (Fig. 5B). The lectin-type agonist promoted the surface accumulation of cytokine receptors by blocking internalization (Fig. 5B) and induced a slow but steady activation of downstream molecules (Fig. 5A) that synergized with natural ligands (Fig. 5C). These data elucidate the previously understudied molecular mechanism of cytokine receptor activation by lectins.

This study presents the first evidence that an exogenous ligand activates MPL via surface glycans on the receptor. Notably, the glycan critical for MPL action is the same one required for activation by the internal ligand CALRmut (*5, 7*). In cells expressing CALRmut, homomultimerized CALRmut engages with MPL bearing immature N-glycans at N117 in the endoplasmic reticulum (ER) to form a 2+2 quadripartite complex of MPL-CALRmut in the Golgi apparatus, and this complex is trafficked to the cell surface for activation (*5, 6, 21*). CALRmut fails to activate MPL expressed on the surface of cells that do not express CALRmut (*10, 11*), which renders the potential of CALRmut as the MPL ligand uncertain and leaves the molecular mechanism of MPL activation ambiguous. The present study clearly demonstrated that the N-glycan at N117 is the bona fide switch for the activation of MPL by homodimerized lectins and ensures the agonistic effect of homomulitimerized CALRmut on MPL.

Our structural insights into ThC led to the discovery of PA-IIL as a ThC-type MPL agonist, which demonstrates the potential of bacterial fucose-binding lectins, including theoretical lectins, as MPL agonists. Structural and biological comparisons of ThC and PA-IIL suggested that the dimerized form and capacity of fucose binding were not sufficient for MPL activation. The position of MPL molecules, which is determined by the positioning of the fucose moiety at N117 of MPL bound by the lectin, plays a crucial role in the degree of receptor activation. Notably, we found that the spatial arrangement of two fucose-binding pockets clearly differed between ThC and PA-IIL. When the core-(1,6) fucose of the sugar chain at N117 of each MPL bound tightly to two ligand binding cores of the lectin, the configuration of the lectin-bound receptor complex directly reflected the spatial relationship of the pocket. Therefore, ThC and PA-IIL-bound activating receptor complexes differ structurally. We propose that this structural difference affects the efficacies of the two lectins. A recent study demonstrated that dimeric antibodies that bridge and dimerize MPL by binding various sites near the canonical ligand-binding domain of MPL activated the receptor in distinctive ways, which realized agonist-based decoupling of HSC self-renewal and differentiation (*22*). Because no structural information on the ‘active’ receptor complex for MPL was known, the mechanistic basis of this phenomenon was elusive. However, our observations, in conjunction with the above study, support the hypothesis that slight structural differences in the extracellular domain of MPL affect receptor activation and signaling.

Our results showed that the ThC-dependent activation of MPL persisted and was associated with sustained expression of MPL on the cell surface. These results were also observed in CALRmut-dependent MPL activation (*5*), suggesting a conserved mechanism of action in lectin-mediated receptor activation. The activation dynamics of MPL are largely controlled by the internalization and recycling of the receptor and the *de novo* biosynthesis of new receptors (*23*). Our results suggested that internalization of ThC-activated receptors was a slow process, as observed in CALRmut-activated receptors, which yielded persistent activation. The strong synergy observed in the ThC/TPO coapplication may be partially ascribed to the sustained signaling of long-lasting cell surface receptors (Fig. S16).

MPL belongs to the class I cytokine receptor family, whose activation relies on the formation of receptor dimers. However, the process of dimer formation is not well understood (*24*). Preformed dimers are likely activated upon conformational alterations (*24, 25*). However, recent single-molecule live cell imaging of MPL expressed on HeLa cells revealed that the population of monomeric receptors surpassed the predimeric form, and the monomer assembled into dimers after interacting with agonists (*26*). Because we observed weak but sustained receptor activation by ThC associated with sustained cell surface expression of the receptor, we propose that a transition state, a *ligand-bound but subtle-activated dimer* in the process of signaling dimer formation, plays an important role (Fig. 5D III). The role of this well-conceivable transition complex formed in the middle of the activation process was previously hidden because this intermediate is short-lived in TPO-activated MPL due to potent ‘native’ interactions between the receptor and ligand (*26*). The strong synergy of ThC with TPO supports this hypothesis because predimerization facilitated by ThC largely reduces entropy during dimer formation. Therefore, ThC stabilizes transition state III, although ThC-bound III eventually shifts to form signaling complex IV, presumably via the aid of intrinsic domain interactions in the transmembrane (TMD) and intracellular domains (Fig. 5D).

Because exogenously applied recombinant CALRmut fails to activate normal MPL (*10, 11*), ThC- and ThC-type fucose-binding lectins were the only probes, and they were excellent for studying the receptor kinetics and dynamics of MPL during N-glycan-mediated activation. The present study is the first report of the structural basis of the sugar-mediated activation of cytokine receptors. MPL-mediated signaling is involved in at least two discrete activation processes in hematopoiesis, hematopoietic progenitor cell differentiation/megakaryocyte formation and HSC self-renewal/maintenance (*1-3*). We propose that fucose-binding lectins are novel tools to control structure to activate the MPL complex and enable fine-tuning of dimerization, internalization, and signaling in conjunction with coapplication with other agonists.

## Supporting information

Watari et al supplemental data

## Acknowledgments

We thank Professor Yasuhiko Kizuka at Gifu University for providing 6-alkynyl fucose and valuable comments. We also thank Dr. Takanori Nakamura at Nissan Chemical Co. Ltd. for providing Ba/F3 cells, the Global Facility Center at Hokkaido University for performing amino acid sequence analysis, Laboratory of Proteomics and Biomolecular Science, Biomedical Research Core Facilities, Juntendo University Graduate School of Medicine for immunoblot data collection, the Teijin Scholarship Foundation, Suntory Foundation for Life Sciences, and Japan Society for the Promotion of Science (JSPS) for the DC2 scholarship to H.W., and the Chuuk State Department of Marine Resources for providing permission to collect sponges.

## Funding

JSPS KAKENHI #19H03040, #21K08405, #19K08848, #21K08376, #21K08424

Ikeda Scientific Co., Ltd.

Grant-in-aid for JSPS fellows #20J11377

Takeda Science Foundation

## Authors’ contributions

This work was conceptualized by H.W., R.S., M.A., and Y.T. H.N., T.O., T.M., Y.K., and H.W. analyzed the protein sequence. H.K., K.O., PJ.F., T.Y., T.O., D.F., and Y.T. overexpressed the proteins and analyzed their crystal structures and biochemical properties. N.M., M.A., and H.W. characterized the biological activities. K.H. and M.H. isolated hypnin. R.S., H.W., Y.T., M.A., and N.M. prepared the manuscript with input from all of the other authors. N.K. supervised the study.

Conceptualization: H.W., R.S., M.A., Y.T., H.K.

Methodology: HW, MA, YT, HK, RS, NM, TM, DMF

Investigation: HW, NM, HK, MA, YT, RS, MH, KH, TaO, YK, PJF, KO, TY

Visualization: RS, HW, NM, MA, YT, HK, TM, ToO

Funding acquisition: R.S., Y.T., M.A., and H.W.

Project administration: R.S., H.W., M.A., and Y.T.

Supervision: N.K., R.S., Y.T., and M.A.

Writing — original draft: R.S., H.W., Y.T., and N.M.

Writing – review & editing: R.S., H.W., Y.T., N.M., and M.A.

## Data and material availability

PDB; 7F9F (nThC), 7F91 (Met substituted rThC), 7F9G (rThC in complex with Ca^2+^ and fucose), 7FBL (rThC in complex with Ca^2+^ and mannose), 7F9J (rThC Q25K in complex with Ca^2+^). All other data are available in the main text or the supplementary materials.

## References

1. A. Nakamura-Ishizu, T. Suda, Multifaceted roles of thrombopoietin in hematopoietic stem cell regulation. Ann N Y Acad Sci 1466, 51–58 (2020).

2. K. Behrens, W. S. Alexander, Cytokine control of megakaryopoiesis. Growth Factors 36, 89–103 (2018).

3. I. S. Hitchcock, K. Kaushansky, Thrombopoietin from beginning to end. British J. Haematol. 165, 259–268 (2014).

4. I. S. Hitchcock, M. Hafer, V. Sangkhae, J. A. Tucker, The thrombopoietin receptor: revisiting the master regulator of platelet production. Platelets 32, 770–778 (2021).

5. N. Masubuchi, M. Araki, Y. Yang, E. Hayashi, M. Imai, Y. Edahiro, Y. Hironaka, Y. Mizukami, Y. Kihara, H. Takei, M. Nudejima, M. Koike, A. Ohsaka, N. Komatsu, Mutant calreticulin interacts with MPL in the secretion pathway for activation on the cell surface. Leukemia 34, 499–509 (2020).

6. C. Pecquet, I. Chachoua, A. Roy, T. Balligand, G. Vertenoeil, E. Leroy, RI. Albu, JP. Defour, H. Nivarthi, E. Hug, E. Xu, Y. O. Amer, C. Mouton, D. Colau, D. Vertommen, M. M. Shwe, C. Marty, I. Plo, W. Vainchenker, R. Kralovics, S. N. Constantinescu, Calreticulin mutants as oncogenic rogue chaperones for TpoR and traffic-defective pathogenic TpoR mutants. Blood 133, 2669–2681 (2019).

7. I. Chachoua, C. Pecquet, M. El-Khoury, H. Nivarthi, RI. Albu, C. Marty, V. Gryshkova, JP. Defour, G. Vertenoeil, A. Ngo, A. Koay, H. Raslova, P. J. Courtoy, M. L. Choong, I. Plo, W. Vainchenker, R. Kralovics, S. N. Constantinescu, Thrombopoietin receptor activation by myeloproliferative neoplasm associated calreticulin mutants. Blood 127, 1325–1335 (2016).

8. C. Marty, C. Pecquet, H. Nivarthi, M. El-Khoury, I. Chachoua, M. Tulliez, JL. Villeval, H. Raslova, R. Kralovics, S. N. Constantinescu, I. Plo, W. Vainchenker, Calreticulin mutants in mice induce an MPL-dependent thrombocytosis with frequent progression to myelofibrosis. Blood 127, 1317–1324 (2016).

9. S. Elf, N. S. Abdelfattah, E. Chen, J. Perales-Patón, E. A. Rosen, A. Ko, F. Peiske, N. Florescu, S. Giannini, O. Wolach, E. A. Morgan, Z. Tothova, JA. Losman, R. K. Schneider, F. Al-Shahrour, A. Mullally, Mutant Calreticulin Requires Both Its Mutant C-terminus and the Thrombopoietin Receptor for Oncogenic Transformation. Cancer Discov. 6, 368–381 (2016).

10. M. Araki, Y. Yang, N. Masubuchi, Y. Hironaka, H. Takei, S. Morishita, Y. Mizukami, S. Kan, S. Shirane, Y. Edahiro, Y. Sunami, A. Ohsaka, N. Komatsu, Activation of the thrombopoietin receptor by mutant calreticulin in CALR-mutant myeloproliferative neoplasms. Blood 127, 1307–1316 (2016).

11. L. Han, L. Han, C. Schubert, J. Köhler, M. Schemionek, S. Isfort, T. H. Brümmendorf, S. Koschmieder, N, Chatain, Calreticulin-mutant proteins induce megakaryocytic signaling to transform hematopoietic cells and undergo accelerated degradation and Golgi-mediated secretion. J. Hematol. Oncol. 9, 45 (2016).

12. H. Watari, H. Nakajima, W. Atsuumi, T. Nakamura, T. Nanya, Y. Ise, R. Sakai, A novel sponge-derived protein thrombocorticin is a new agonist for thrombopoietin receptor. Comp. Biochem. Physiol. C Toxicol. Pharmacol. 221, 82–88 (2019).

13. R. Loris, D. Tielker, K. E. Jaeger, L. Wyns, Structural basis of carbohydrate recognition by the lectin LecB from Pseudomonas aeruginosa. J. Mol. Biol. 331, 861–870 (2003).

14. E. Mitchell, C. Houles, D. Sudakevitz, M. Wimmerova, C. Gautier, S. Perez, A. M. Wu, N. Gilboa-Garber, A. Imberty, Structural basis for oligosaccharide-mediated adhesion of Pseudomonas aeruginosa in the lungs of cystic fibrosis patients. Nat. Struct. Biol. 9, 918–921 (2002).

15. C. Sabin, C. Sabin, E. P. Mitchell, M. Pokorná, C. Gautier, JP. Utille, M. Wimmerová, A. Imberty, Binding of different monosaccharides by lectin PA-IIL from Pseudomonas aeruginosa: thermodynamics data correlated with X-ray structures. FEBS Lett. 580, 982–987 (2006).

16. Y. Kizuka, M. Nakano, Y. Yamaguchi, K. Nakajima, R. Oka, K. Sato, CT. Ren, TL. Hsu, CH. Wong, N. Taniguchi, An Alkynyl-fucose halts hepatoma cell migration and invasion by Inhibiting GDP-Fucose-Synthesizing Enzyme FX, TSTA3. Cell Chem. Biol. 24, 1467–1478 e1465 (2017).

17. J. Li, H. C. Hsu, J. D. Mountz, J. G. Allen, Unmasking Fucosylation: from Cell Adhesion to Immune System Regulation and Diseases. Cell. Chem. Biol. 25, 499–512 (2018).

18. S. Okuyama et al., Strict binding specificity of small-sized lectins from the red alga Hypnea japonica for core (alpha1-6) fucosylated N-glycans. Biosci. Biotechnol. Biochem. 73, 912–920 (2009).

19. R. I. Albu, S. N. Constantinescu, Extracellular domain N-glycosylation controls human thrombopoietin receptor cell surface levels. Front. Endocrinol. 2, 71 (2011).

20. S. Elf, N. S. Abdelfattah, A. J. Baral, D. Beeson, J. F. Rivera, A. Ko, N. Florescu, G. Birrane, E. Chen, A. Mullally, Defining the requirements for the pathogenic interaction between mutant calreticulin and MPL in MPN. Blood 131, 782–786 (2018).

21. M. Araki, Y. Yang, M. Imai, Y. Mizukami, Y. Kihara, Y. Sunami, N. Masubuchi, Y. Edahiro, Y. Hironaka, S. Osaga, A. Ohsaka, N. Komatsu, Homomultimerization of mutant calreticulin is a prerequisite for MPL binding and activation. Leukemia 33, 122–131 (2019).

22. L. Cui, I. Moraga, T. Lerbs, C. V. Neste, S. Wilmes, N. Tsutsumi, A. C. Trotman-Grant, M. Gakovic, S. Andrews, J. Gotlib, S. Darmanis, M. Enge, S. Quake, I. S. Hitchcock, J. Piehler, K. C. Garcia, G. Wernig, Tuning MPL signaling to influence hematopoietic stem cell differentiation and inhibit essential thrombocythemia progenitors. Proc. Natl. Acad. Sci. U.S.A. 118, (2021).

23. D. D. Dahlen, V. C. Broudy, J. G. Drachman, Internalization of the thrombopoietin receptor is regulated by 2 cytoplasmic motifs. Blood 102, 102–108 (2003).

24. L. N. Varghese, J. P. Defour, C. Pecquet, S. N. Constantinescu, The Thrombopoietin Receptor: Structural Basis of Traffic and Activation by Ligand, Mutations, Agonists, and Mutated Calreticulin. Front. Endocrinol. 8, 59 (2017).

25. A. J. Brooks, M. J. Waters, The growth hormone receptor: mechanism of activation and clinical implications. Nat. Rev. Endocrinol. 6, 515–525 (2010).

26. S. Wilmes, M. Hafer, J. Vuorio, J. A. Tucker, H. Winkelmann, S. Löchte, T. A. Stanly, K. D. P. Prieto, C. Poojari, V. Sharma, C. P. Richter, R. Kurre, S. R. Hubbard, K. C. Garcia, I. Moraga, I. Vattulainen, I. S. Hitchcock, J. Piehler, Mechanism of homodimeric cytokine receptor activation and dysregulation by oncogenic mutations. Science 367, 643–652 (2020).

27. T. Nakamura, Y. Miyakawa, A. Miyamura, A. Yamane, H. Suzuki, M. Ito, Y. Ohnishi, N. Ishiwata, Y. Ikeda, N. Tsuruzoe, A novel nonpeptidyl human c-Mpl activator stimulates human megakaryopoiesis and thrombopoiesis. Blood 107, 4300–4307 (2006).

28. T. Masuda, M. Tomita, Y. Ishihama, Phase transfer surfactant-aided trypsin digestion for membrane proteome analysis. J. Proteome Res. 7, 731–740 (2008).

29. J. Rappsilber, M. Mann, Y. Ishihama, Protocol for micro-purification, enrichment, pre-fractionation and storage of peptides for proteomics using StageTips. Nat. Protoc. 2, 1896–1906 (2007).

30. Z. Otwinowski, W. Minor, Processing of X-ray Diffraction Data Collected in Oscillation Mode. Methods in Enzymol. 276, 307–326 (1997)

31. W. Kabsch, Xds. Acta. Crystallogr. D Biol. Crystallogr. 66, 125–132 (2010)

32. P. Thomas, R.S. Thomas, HKL2MAP: a graphical user interface for macromolecular phasing with SHELX programs. J.Appl. Crystallogr. 37, 843–844 (2004)

33. T.C. Terwilliger, P.D. Adams, R.J. Read, A.J. McCoy, N.W. Moriarty, R.W. Grosse-Kunstleve, P.V. Afonine, P.H. Zwart, and L.W. Hung, Decision-making in structure solution using Bayesian estimates of map quality: the PHENIX AutoSol wizard. Acta Crystallogr. D Biol. Crystallogr. 65, 582–601 (2009).

34. A.J. McCoy, R.W. Grosse-Kunstleve, P.D. Adams, M.D. Winn, L.C. Storoni, and R.J. Read. Phaser crystallographic software. J. Appl. Crystallogr. 40, 658–674 (2007).

35. P.V. Afonine, R.W. Grosse-Kunstleve, N. Echols, J.J. Headd, N.W. Moriarty, M. Mustyakimov, T.C. Terwilliger, A. Urzhumtsev, P. H. Zwart and P. D. Adams, Towards automated crystallographic structure refinement with phenix.refine. Acta Crystallogr D Biol. Crystallogr. 68, 352–367 (2012).

