## Supplementary material for "Hidden pathway for cytokine receptor activation: Structural insights into a marine sponge-derived lectin that activates the thrombopoietin receptor via recognition of the fucose moiety": Watari et al supplemental data

### Materials and Methods

#### Affinity purification of native ThC

The sponge specimen used here was collected in Chuuk state of Federated States of Micronesia in 2009 under permission of Department of Marine Resources, Chuuk State FSM, and was extracted as described previously (*12*). A sponge aqueous extract was treated with acidic buffer (pH 3.0) to give ThC-enriched extract. The extract was applied on a 1 mL of Sepharose-fucose affinity gel (EY Laboratories, Inc., San Mateo, CA). The column was eluted first with 50 mM Tris-HCl buffer and then by fucose. The fucose eluent was dialyzed to give purified protein.

#### Cell culture and proliferation assay

Cell proliferation activities of ThC were carried out as described previously (*12*). Briefly, murine interleukin-3 dependent pro-B cell line Ba/F3 expressing human *mpl* (Ba/F3-HuMpl cells) (*27*) were pre-cultured for 4 days, and then harvested by centrifugation at 1,000 rpm for 3 min. After washing with PBS (-), the collected cells were resuspended in RPMI-1640 medium containing 10% FBS at a cell density of 6.0 x $10^4$ cells/mL. A 90 µL aliquot of the cell resuspension were transferred to 96 well plate. In the presence of various concentrations of ThC, or recombinant ThCs for 10 µL, cells were cultivated at 37°C under 5% $CO_2$ concentration. As negative and positive controls, PBS and TPO (PeproTech, 300-18, NJ, USA) were used. After 4 days cultivation, cell proliferation was measured with cell counting kit (Dojindo, Kumamoto, Japan). A 10 µL aliquot of cell counting kit was added to each well. After incubation for 2 hours, absorption at 450 nm ($Abs_{450}$) was recorded with microplate reader. For 6-Alkynyl-

Fucose assay, Ba/F3-HuMpl cells were pre-treated with 6-Alk-Fuc for 6 hours, then added ThC or TPO and cultured for 4 days.

### Preparation of samples for the cell proliferation assay

ConA (SIGMA), hypnin (from *Hypnea japonica*), PA-IIL (Fujifilm-Wako), BC2LC-CTD (recombinant) were dissolve in PBS (-). 6-alkynyl fucose (Peptide Institute, INC., Japan) was dissolved at 100 mM in DMSO and diluted with PBS (-) to each concentration. Sugar solutions were prepared with PBS (-) except N-acetylneuraminic acid (NANA). NANA was suspended in PBS (-) and neutralized with aqueous NaOH to pH7.0. All the reagents were filter-sterilized with a 0.2 μm filter prior to use.

### Amino acid sequence determination

Draft amino acid Edman degradation was performed using ThC purified by SDS-PAGE. The gel was electroblotted on a PVDF membrane and the band for ThC was cutout for *N*-terminal amino acid sequence using automated sequencer (Procise 492HT, Parkin Elmer). Internal amino acid sequence was obtained by digesting purified ThC with either trypsin (Fujifilm-Wako), chymotrypsin (Fujifilm-Wako) or V8 protease (Fujifilm-Wako). Each digest was separated by an HPLC using a reversed-phase column (VYDAC protein&peptide C18) with a gradient (0-50%) of 0.1% aqueous TFA and acetonitrile. Each of the peptide fragments was subjected to de novo sequence analysis using MALDI-TOF MS/MS and to Edman degradation. Deduced amino acid sequences were mapped to give draft amino acid sequence of ThC. Next, entire amino acid sequence was confirmed by mass-spectrometry as follows. A drop (4.5 µL) of crystallization supernatant from X-ray analysis experiment containing 7.0 µg of the native ThC was mixed with 80 µL of acetone, and then centrifuged 19,000 x g for 15 min at 4 ºC. The precipitate was resuspended with 80 µL of acetone. After centrifugation of 19,000 x g for 15 min at 4 ºC, the precipitate was further washed as above and then was air-dried. The dried sample was resuspended to 40 µL of 1× phase transfer surfactant (PTS) (*28*). A 20 µL of resuspended sample was incubated with 2 µL of 200 mM Bond-Breaker TCEP solution (Thermo Fisher Scientific MA, USA) to cleave disulfide bond for 30 min at 50ºC. The reduced thiol was alkylated by 2 µL of 375 mM 2-iodoacetamide for 30 min at room temperature in the dark. After alkylation, an excess amount of 2-iodoacetamide was reacted with 2 µL of 400 mM L-Cys for 10 min at room temperature in the dark. The alkylated sample was digested by either 200 ng of trypsin (Promega, USA) with 200 ng of Lys-C (Fujifilm-Wako) at 37ºC overnight or 200 ng of chymotrypsin (Promega) with 10 mM $CaCl_2$ at 25ºC overnight. A total of 30 µL of the digestion sample was precipitated by the addition of 45 µL of 1.7% trifluoroacetic acid (TFA). After centrifugation at 19,000 x g for 15 min at 4 ºC, the supernatant was purified using Stage-Tip described previously (*29*). Peptide fragments were eluted from Stage-Tip using 70% acetonitrile and 0.1% TFA, and the elution was freeze-dried. 2.2 µg of recombinant ThC (rThC Q25) dissolved in 20 µL of PTS was also digested followed by the protocol described above. The peptide fragments were dissolved with 10 µL of 0.1% TFA, and the peptides of the native ThC and rThC Q25 were analyzed with a quadrupole Orbitrap benchtop mass spectrometer (Q-Exactive, Thermo Fisher Scientific) equipped with a Nanospace SI-2 HPLC system (Osaka Soda Co., Ltd., Osaka, Japan), respectively. The column temperature was maintained at 45 °C. The flow rate of the mobile phase was 200 µL/ min, and mobile phase A consisted of 0.05% formic acid (FA) and mobile phase B consisted of 0.05% FA/90%

acetonitrile. The mobile phase gradient was programmed as follows: 0% B (0-2 min), 0-35% B (2-12 min), 35-55% B (12-15 min), 55-80% B (15-16 min), 80% B (16-18 min), 80-0% B (18-18.5 min), and 0% B (18.5-20 min). MS1 spectra were collected in the scan range of 350−1,200 m/z at 70,000 resolution at 200 m/z to hit an AGC target of $1\times10^6$ with an injection time of 200 ms. AGC target value for fragment spectra was set at $1\times10^5$, and the intensity threshold was kept at $3.3\times10^4$. Isolation width was set at 2.4 m/z, the 12 most intense ions were fragmented in a data-dependent mode by collision-induced dissociation with a normalized collision energy of 27.

The amino acid sequences of draft sequence and rThC Q25 were added into the UniProt sequence database (release 31st July 2019, entry 557,016, all species, reviewed). MS file was searched against the database using Proteome Discoverer 1.4 software (Thermo Fisher Scientific) and PEAKS Studio Version X (Bioinformatics Solutions. Waterloo, Canada). The setting parameters were as follows: enzyme, trypsin (semi) or chymotrypsin (semi); maximum missed cleavage sites, 2 (Proteome Discoverer) or 4 (PEAKS); precursor mass tolerance, 6 ppm; fragment mass tolerance, 0.02 Da; fixed modification, cysteine carbamidomethylation. The peptides' identification was filtered to a false discovery rate of less than 1%.

### Over expression and purification of recombinant ThC (rThC), rThC mutants, and BC2LC-CTD

Expression vector of recombinant ThC (rThC) and BC2LC-CTD was constructed by cloning a synthesized DNA fragment corresponding to the amino acid sequence of ThC determined by mass spectrometry and C-terminal domain of BC2LC into NdeI/XhoI site of a modified pET26 and pET28 vector, respectively. The 6 × His tag was attached to the N-terminus of ThC. Expression vector of rThC mutants was constructed by inverse PCR method using the expression vector of rThC as a template.

*Escherichia coli* strain BL21(DE3) harboring the expression vector of the desired protein was cultivated in LB medium at 37 °C with shaking 120 rpm. When $OD_{600}$ reaches to 0.6, isopropyl-β-D-thiogalactopyranoside (IPTG) was added to the medium at the final concentration of 0.2 mM to induce expression of the rThC, followed by further incubation at 25 °C for overnight.

Cells were harvested by centrifugation at 4,000 x g for 30 min. The collected cell was suspended in buffer A composed of 20 mM HEPES-NaOH (pH 8.0), and 200 mM NaCl, and then disrupted with UD-211 ultrasonic disruptor (TOMY SEIKO, Japan). After centrifugation at 40,000 x g for 30min, the supernatant was loaded onto a 1 mL column of Ni Sepharose (GE Healthcare, Sweden). After washing with sonication buffer, the bound protein was eluted using a concentration gradient of imidazole in the sonication buffer. Fractions containing purified rThC were further purified by size exclusion chromatography using HiLoad 26/600 Superdex 75pg (GE Healthcare) preequilibrated with the buffer A. Fractions containing rThC were collected and used for further experiments. SeMet substituted rThC was expressed and purified by the same method as rThC except using SeMet substituted M9 medium instead of LB medium.

### Isothermal Titration Calorimetry (ITC)

ITC measurement was carried out with an iTC200 (GE healthcare). The cell was filled with approximately 100 µM rThC, 100 µM BC2LC-CTD, or 35 µM PA-IIL, and the syringe was filled with 1.5 μM fucose or mannose. The protein was injected 18 times in portion of 2 μL over 120 sec. The data were analyzed with the program ORIGIN.

**Carbohydrates binding assay**

Carbohydrate binding specificity of rThC was analyzed with carbohydrate Gel Kit#1 (EY Laboratories). Binding to fucose, mannose, lactose, N-acetylglucosamine, and N-acetylgalactosamine was evaluated with resin in which each carbohydrate was immobilized. A 0.5 mL aliquot of 0.1 mg/mL of the purified rThC was loaded on 0.1 mL of the resin immobilizing each carbohydrate. After washing with 0.5-mL of buffer, the bound rThC was competitively eluted by elution buffer containing 0.2 M of carbohydrate immobilized on the resin. rThC specifically bound to the carbohydrate was evaluated by SDS-PAGE.

**Crystallization, X-ray diffraction data collection, and structure determination**

For crystallization, purified proteins concentrated up to approx. 6 mg/mL were used. Crystals of nThC was grown from a buffer composed of 0.1 M Tris-HCl (pH 8.5), 0.2 M $MgCl_2$, 30% (w/v) PEG 4000. The diffraction dataset of the nThC was collected at Advanced Photon Source (IL, USA). The diffraction data of nThC were processed with the program HKL2000 *(30)*.

For phasing of the nThC, crystal structure of SeMet substituted rThC (SeMet-rThC) were determined. Crystals of SeMet-rThC were grown from a buffer composed of 100 mM sodium acetate (pH 3.3~5.5), 20% PEG3350~6000, and 20% PEG400. rThC Q25K was crystallized in the presence of 5 mM $CaCl_2$, as biochemical analysis revealed that rThC requires $Ca^{2+}$ ion for its carbohydrate binding activity. Crystals of rThC Q25K in the presence of 5 mM $CaCl_2$ were grown from a buffer composed of 100 mM sodium acetate (pH 3.3~5.5), 20% PEG3350~6000, and 20% PEG400. Crystals of rThC in complex with fucose or mannose were obtained by co-crystallization, in which 5 mM of fucose or mannose was added to the purified rThC solution. X-ray diffraction experiments were carried out in Photon Factory (Tsukuba, Japan) and SPring-8 (Harima, Japan). Diffraction data of SeMet substituted rThC, rThC in the presence of $CaCl_2$, rThC in complex with fucose, and rThC in complex with mannose were collected in Photon Factory. The diffraction data of rThC were processed with the program XDS *(31)*. The statistics of data collection are summarized in Table S2.

Crystal structure of SeMet-rThC was determined by Se-SAD method. The sites of Se were determined using the program HKL2MAP *(32)*. Phasing and model building were carried out using phenix.autosol *(33)*. Crystal structure of nThC was determined by molecular replacement method using the program phenix.mr *(34)* with the structure of SeMet-rThC as search probe. Crystal structures of rThC and its complexes with $Ca^{2+}$ ion, mannose, and fucose were determined by molecular replacement method with the structure of nThC as a search probe. Anomalous difference Fourier map was calculated using phenix.maps. Structure refinement was carried out using phenix.refine *(35)*.

#### STAT5 Reporter assay

For the expression MPL with an amino acid substitution from asparagine (N) to Glutamine (Q) on potential N-glycosylation site at N117, 178, 298, and 358, cDNAs were created by PCR mutagenesis (primers listed in supplementary Table S2) and subcloned into the pcDNA3.1 vector (Life Technologies). V5-tagged mutant MPL and untagged wild-type MPL were used for reporter assay. All plasmids constructed were verified by sequencing before use. Reporter assay was performed as described previously (*21*) with following modifications. The reporter activity was measured 24 hours after the transfection and 19 hours after the addition of ThC or TPO.

#### Immunoblot

All immunoblot analyses were performed as described previously (*10*). The following primary antibodies purchased from Cell Signaling were additionally used in this study: anti-STAT5(#94205).

#### Measurement the levels of MPL on cell surface

Cell surface MPL was isolated and detected as described previously (*5*) with following modifications. Ba/F3-HuMpl cells were pre-cultured with the medium in the absence of agonist overnight, and then incubated with the media in the presence or absence of ThC or TPO at 37°C in a humidified incubator with 5% $CO_2$. For the examination of synergistic effect of ThC and TPO, the pre-culture was performed with the media in the absence of agonist overnight and in the presence of ThC for 2 hours before the addition of TPO.

#### Statistical analysis

Data were obtained more than triplicate at each datapoint representing mean value and error bars ±SD. Concentration response curves are generated using GraphPad Prism

8.0.3. $EC_{50}$ values with 95% confident intervals were obtained by dose response curve fittings with nonlinear regression curve fit with four parameters.

### Table S1. Affinities and thermodynamic parameters of binding

| protein | ligand | 5mM $CaCl_2$ | $K_D$ (μM)[a] | ΔH (kcal $mol^{-1}$)[b] | ΔS (cal $mol^{-1}$ $K^{-1}$)[b] | $K_A$ ($M^{-1}$ x $10^4$)[b] | $K_D$ in the previous report (μM) |
|---|---|---|---|---|---|---|---|
| rThC | fucose | + | 4.72 ± 0.25 | -8.14 | -2.94 | 21.2 ± 1.12 | |
| | | - | NB | NB | NB | NB | |
| | mannose | + | 66.2 ± 15.3 | -0.76 | 16.6 | 1.51 ± 0.35 | |
| | | - | NB | NB | NB | NB | |
| Q25K | fucose | + | NB | NB | NB | NB | |
| G132 | fucose | + | NB | NB | NB | NB | |
| BC2LC-CTD | fucose | + | NB | NB | NB | NB | |
| | mannose | + | 17.7 ± 0.32 | -10.0 | -11.9 | 5.65 ± 0.42 | 37.4[c] |
| PA-IIL | fucose | + | 2.78 ± 0.30 | -6.46 | 3.75 | 35.9 ± 3.90 | 2.9 ± 0.03[d] |
| | mannose | + | 78.7 ± 23.6 | -9.46 | -12.9 | 1.27 ± 0.38 | (Me-α-Man: 71 ± 3)[d] |

Not bound is indicated by "NB".

*a* $K_D = 1/K_A$

*b* $RT \ln K_A = \Delta H - T\Delta S$

*c* https://doi.org/10.1371/journal.ppat.1002238

*d* https://febs.onlinelibrary.wiley.com/doi/full/10.1016/j.febslet.2006.01.030

### Table S2. Data collection and refinement statistics

| | | Native ThC | SeMet Substituted rThC | rThC in complex with $Ca^{2+}$ and fucose | rThC in complex with $Ca^{2+}$ and mannose | rThC Q25K in complex with $Ca^{2+}$ |
|---|---|---|---|---|---|---|
| PDB ID | | 7F9F | 7F91 | 7F9G | 7FBL | 7F9J |
| Beamline | | APS BL 21-ID-F | Photon Factory BL-1A | Photon Factory NE3A | Photon Factory NE3A | Photon Factory NE3A |
| Data collection | | | | | | |
| Space group | | $P\,2_1$ | $P\,2_12_12_1$ | $P\,2_12_12_1$ | $P\,2_12_12_1$ | $P\,2_12_12_1$ |
| Cell dimensions | | | | | | |
| | $a, b, c$ (Å) | 42.46, 92.44, 58.54 | 41.80, 45.00, 109.91 | 41.52, 44.72, 108.58 | 41.57, 44.98, 109.26 | 41.66, 44.88, 109.56 |
| | $\alpha, \beta, \gamma$ (°) | 90, 102.8, 90 | 90, 90, 90 | 90, 90, 90 | 90, 90, 90 | 90, 90, 90 |
| Resolution (Å) $^a$ | | 30.47 - 1.41 (1.43 - 1.41) | 41.64 - 1.40 (1.49 - 1.40) | 41.35 -1.33 (1.41 - 1.33) | 41.59 - 1.42 (1.50 - 1.42) | 38.94 - 1.10 (1.17 - 1.10) |
| $R_{sym}$ (%) $^a$ | | 9.9 (64.2) | 9.6 (66.3) | 5.7 (37.7) | 7.7 (50.3) | 5.3 (65.8) |
| $<I/\sigma(I)>$ $^a$ | | 25.15 (2.17) | 11.28 (2.14) | 26.05 (5.64) | 22.27 (4.64) | 21.83 (2.70) |
| Completeness (%) $^a$ | | 99.9 (99.7) | 99.9 (99.6) | 99.8 (98.7) | 99.7 (98.7) | 99.4 (96.6) |
| Redundancy $^a$ | | 5.7 (4.4) | 7.04 (7.07) | 9.60 (9.48) | 9.51 (9.31) | 9.19 (7.57) |
| Wilson B-factor (Å$^2$) | | 10.3 | 13.6 | 9.1 | 9.6 | 10.9 |
| Refinement | | | | | | |
| No. of reflections used in refinement | | 81704 | 41580 | 47088 | 39532 | 83639 |
| No. of reflections used for $R_{free}$ | | 1950 | 2080 | 1642 | 1369 | 2900 |
| $R_{work}$ / $R_{free}$ | | 0.169 / 0.194 | 0.175 / 0.200 | 0.155 / 0.168 | 0.162 / 0.182 | 0.185 / 0.197 |
| No. atoms | | | | | | |
| | Protein | 3854 | 1940 | 2009 | 2017 | 2023 |
| | Ligand | 2 | 0 | 26 | 27 | 2 |
| | Water | 799 | 247 | 323 | 317 | 332 |
| R.m.s deviations | | | | | | |
| | Bond lengths (Å) | 0.005 | 0.005 | 0.005 | 0.007 | 0.005 |
| | Bond angles (°) | 0.89 | 0.89 | 0.93 | 0.97 | 0.93 |
| Average B-factor (Å$^2$) | | 15.4 | 17.0 | 11.6 | 11.1 | 14.3 |
| | Protein | 13.5 | 16.1 | 10.0 | 9.7 | 12.8 |
| | ligand | 7.9 | - | 14.7 | 7.4 | 18.1 |
| | Water | 24.9 | 24.3 | 21.6 | 20.4 | 23.2 |
| Ramachandran plot | | | | | | |
| | Favored (%) | 97.67 | 97.67 | 97.67 | 97.28 | 98.05 |
| | Allowed (%) | 2.33 | 2.33 | 2.33 | 2.33 | 1.95 |
| | Disallowed (%) | 0 | 0 | 0 | 0.39 | 0 |

*a* The values in parentheses refer to data in the highest resolution shell.

**Table S3. Primers used for the construction of MPL derivatives**

| Primer name | Primer sequence |
|---|---|
| MPL N117Q_Fw | 5'-GTGTTCCTACAGCAGACTCGGACTCAGCGAGTCC-3' |
| MPL N117Q_Rev | 5'-CGAGTCTGCTGTAGGAACACATTCTTCACCCAG-3' |
| MPL N178Q_Fw | 5'-GATCCCAAGCAGTCCACTGGTCCCACGGTCATACAG-3' |
| MPL N178Q_Rev | 5'-CCAGTGGACTGCTTGGGATCTCTGGGGCCATAGC-3' |
| MPL N298Q_Fw | 5'-GACCTGAAGCAGGTTACCTGTCAATGGCAGCAAC-3' |
| MPL N298Q_Rev | 5'-CAGGTAACCTGCTTCAGGTCCAAGGTAAAGCATTGC-3' |
| MPL N358Q_Fw | 5'-CAAGTCACGACAGGACAGCATTATTCACATCCTTG-3' |
| MPL N358Q_Rev | 5'-GCTGTCCTGTCGTGACTTGAAGTGGCAGCGAGAG-3' |

.

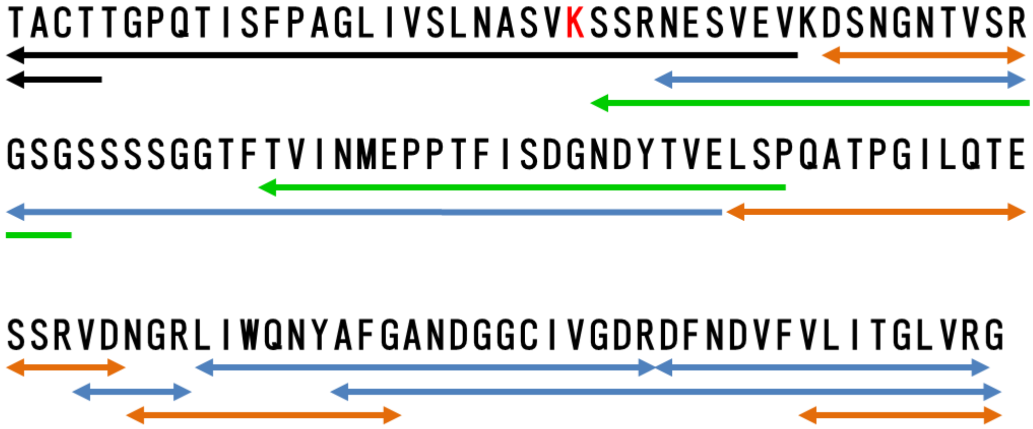

**Fig. S1. Draft amino acid sequence of nThC**

Amino acid sequence of native ThC was first determined on the basis of Edman degradation (black line), and MALDI-TOF MS/MS *de novo* sequecing data of chymotrypsin (green), trypsin, (blue), and V8 protease (orange) digests. The 25$^{th}$ residue was assignes to by lysine (red letter).

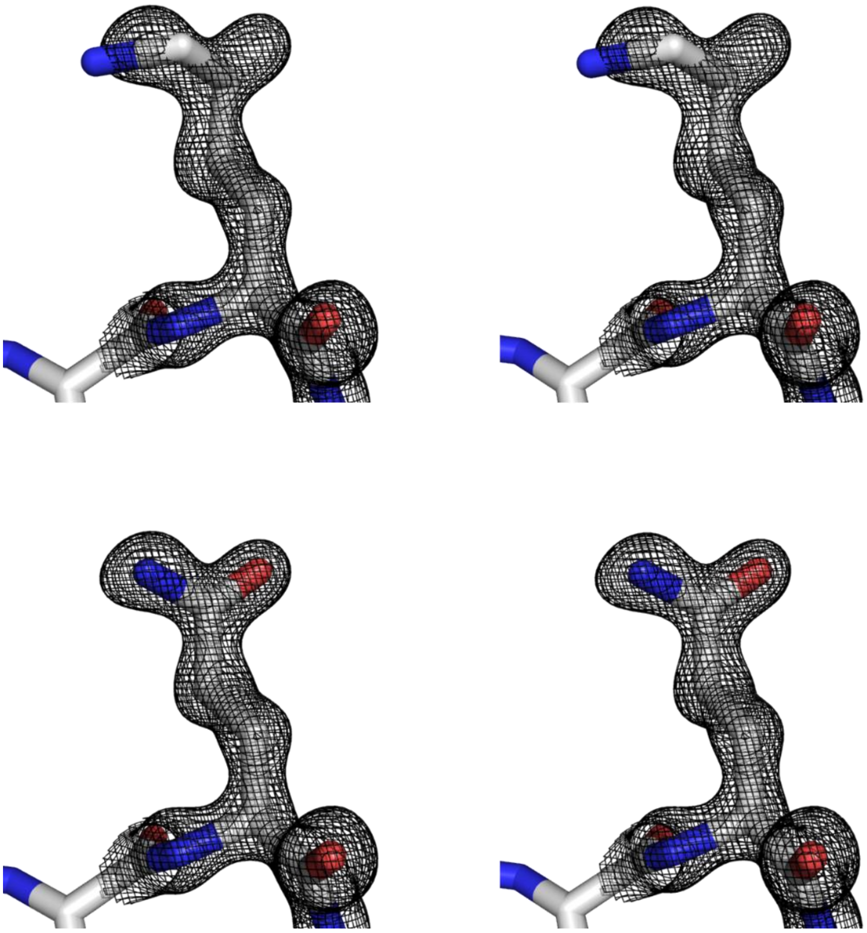

**Fig. S2. Electron density of the 25th residue (stereo view)**

Atomic models of Lys (top) and Gln (bottom) are in the electron density of nThC. Gln fits well (bottom) whereas Lys does not with the electron density (top).

Fig.S. 3A

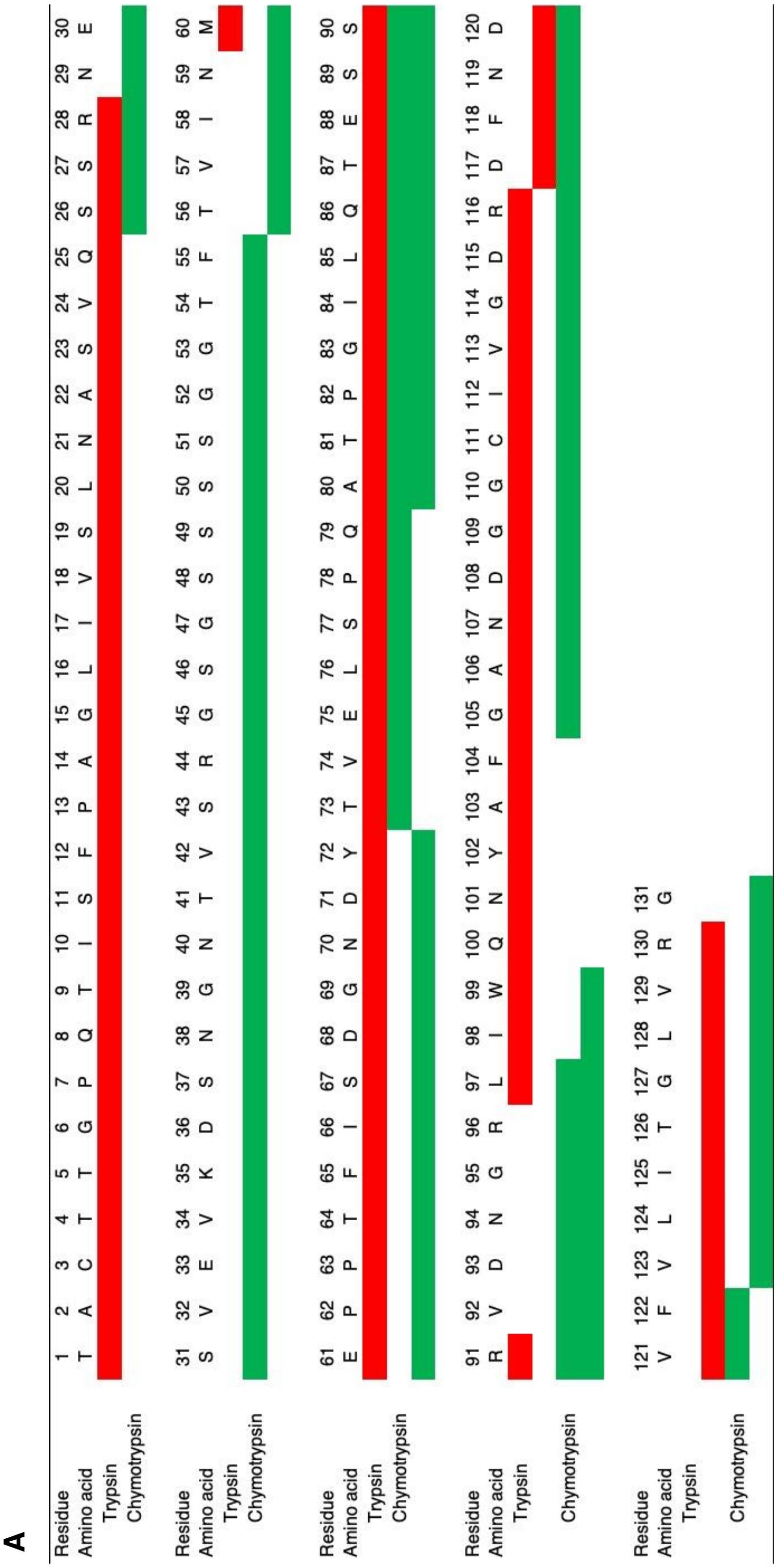

Fig. S3. B-1

**B** TAcTTGPQTISFPAGLIVSLNASVQSSR (res 1-28)

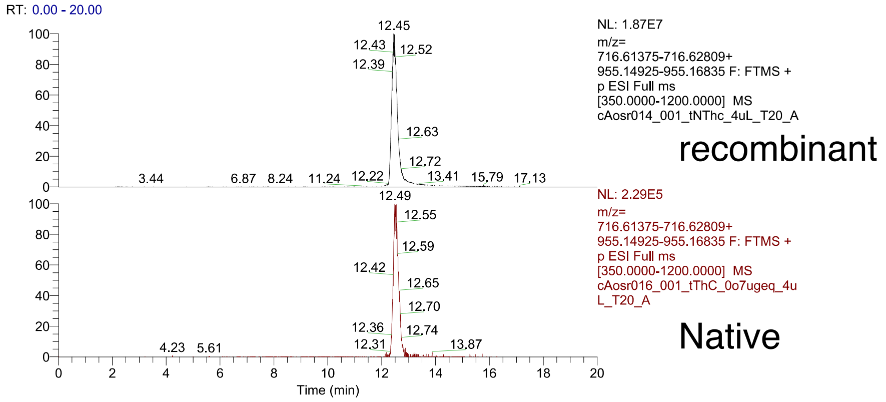

charge 4 (theoretical m/z=716.62092)
recombinant m/z=716.62140, RT=12.62 Δppm = 0.63 (PD)
Native

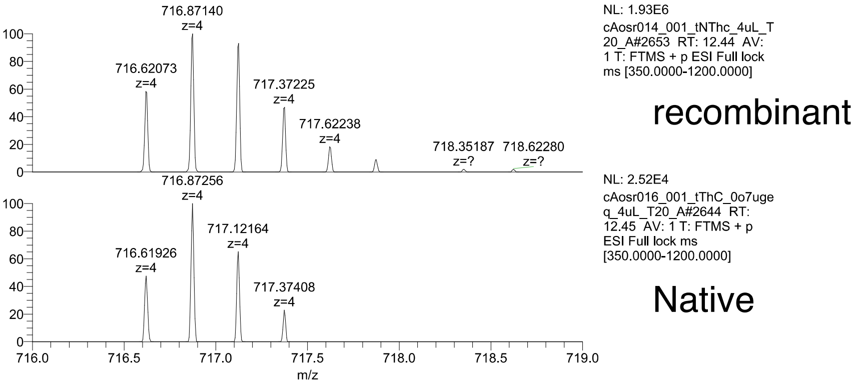

charge 3 (theoretical m/z=955.15880)
recombinant
Native m/z=955.15845, RT=12.38, Δppm=-0.41 (PD)

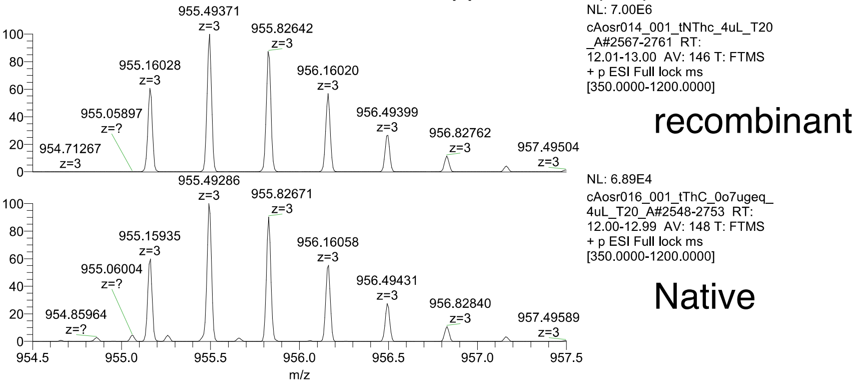

Fig. S3. B-2

### SSRNESVEVKDSNGNTVSRGSGSSSSGGTF (res 26-55)

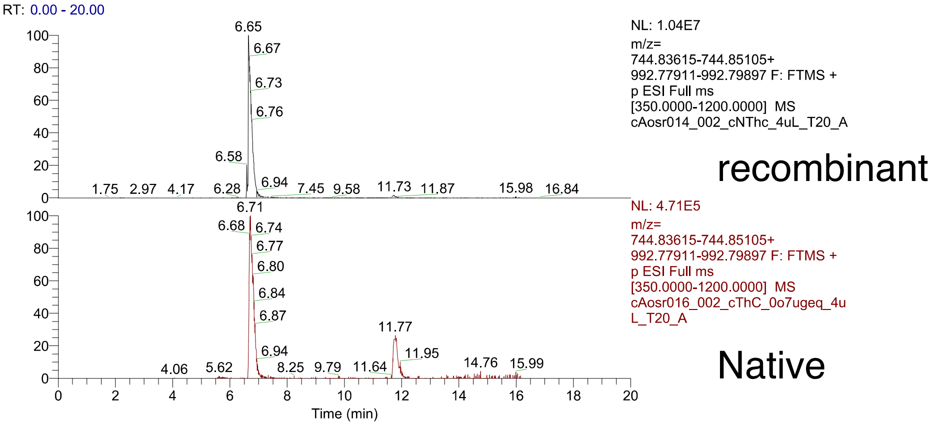

charge 3 (theoretical m/z=992.78904)
recombinant m/z=992.78967, RT=6.59, Δppm=0.60 (PD)
Native

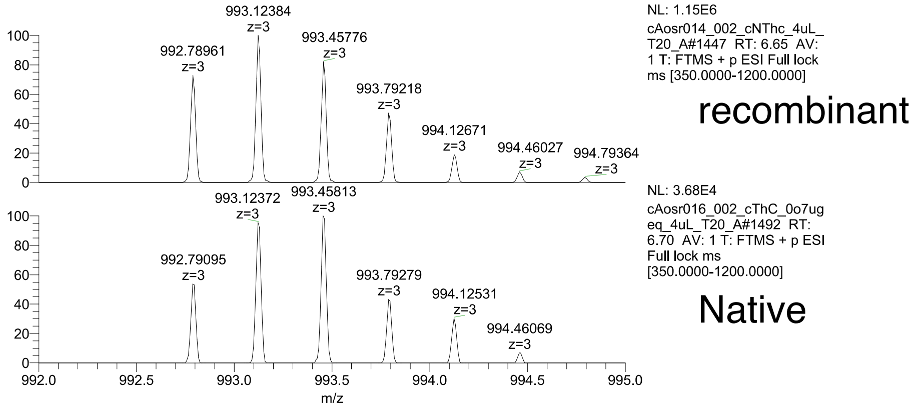

charge 4 (theoretical m/z=744.84360)
recombinant m/z=744.8442, RT=6.57, Δppm=0.8 (PEAKS)
Native m/z=744.8444, RT=6.71, Δppm=1.0 (PEAKS)

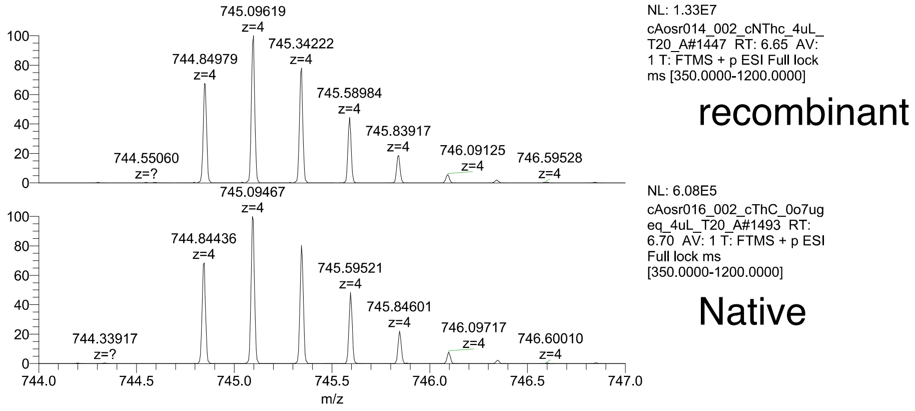

Fig. S3. B-3

### TVINMEPPTFISDGNDY (res 56-72)

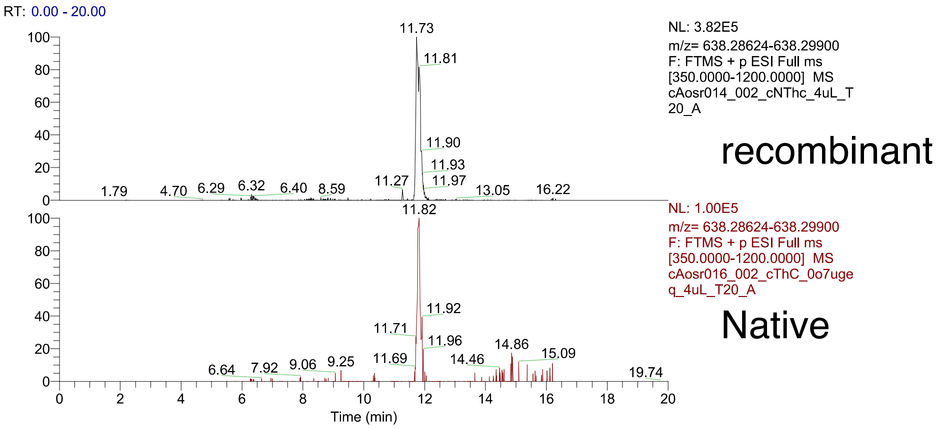

charge 3 (theoretical m/z=638.29262)
recombinant m/z=638.2927, RT=11.71, Δppm=0.2 (PEAKS)
Native m/z=638.2935, RT=11.80, Δppm=1.4 (PEAKS)

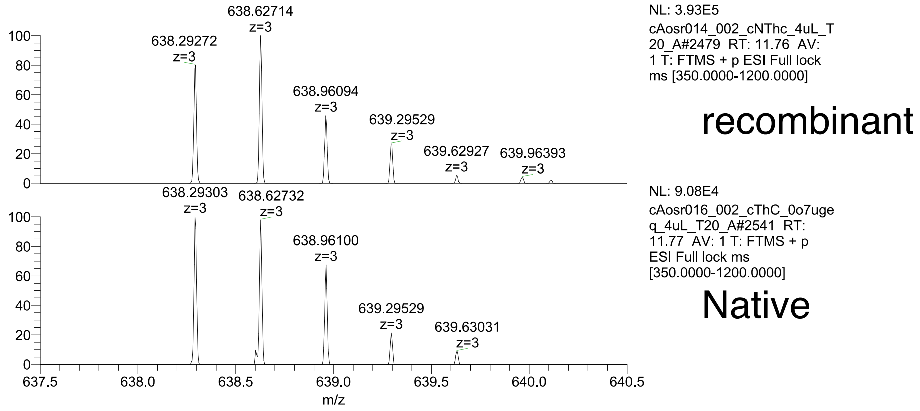

Fig. S3. B-4

MEPPTFISDGNDYTVELSPQATPGILQTESSR (res 60-91)

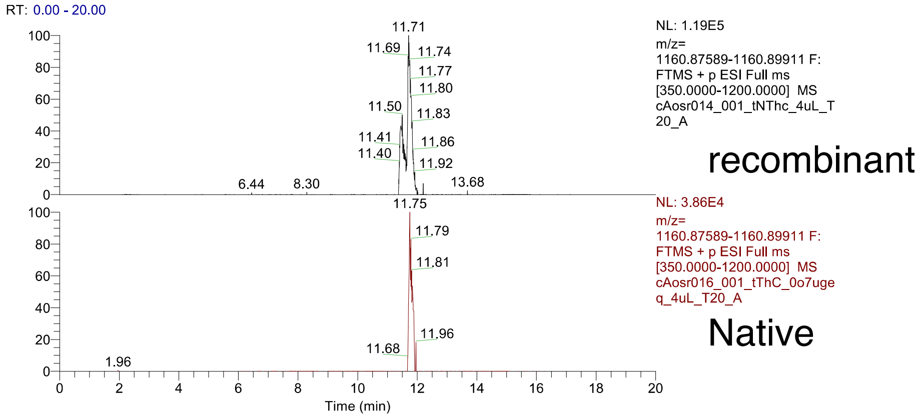

charge 3 (theoretical m/z=1160.88736)
recombinant
Native m/z=1160.8875, RT=11.75, Δppm=0.1 (PEAKS)

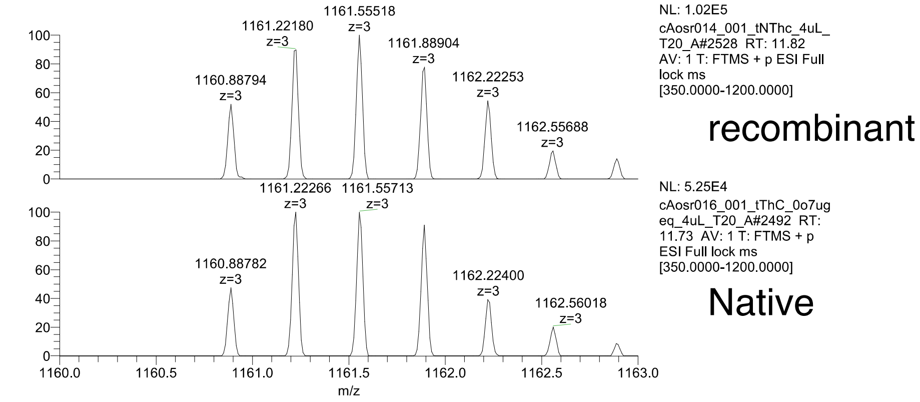

Fig. S3. B-5

### TVELSPQATPGILQTESSRVDNGRL (res 73-97)

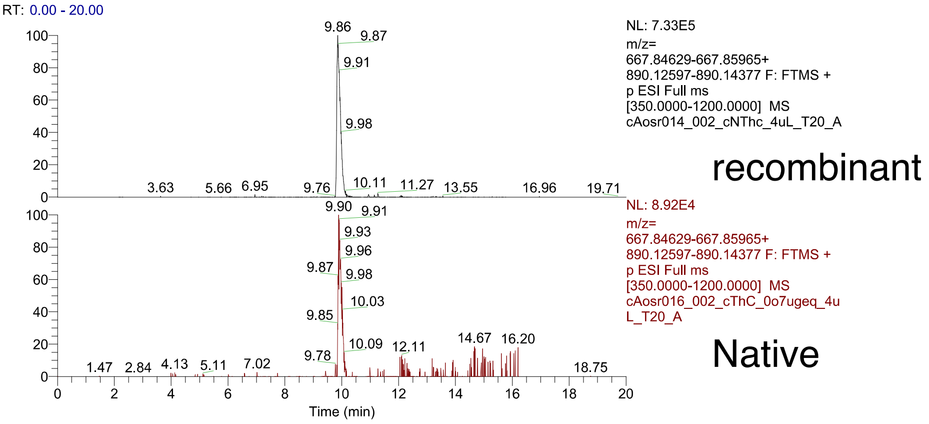

charge 4 (theoretical m/z=667.85297)
recombinant m/z=667.85278, RT=9.83, Δppm=-0.33 (PD)
Native

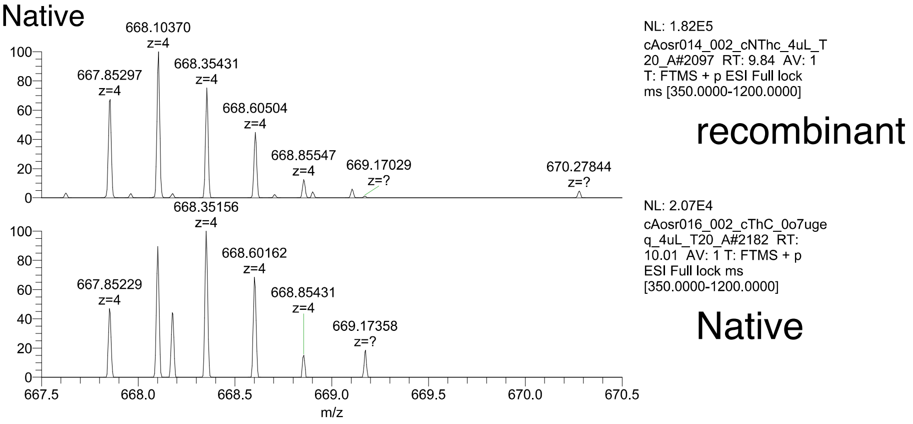

charge 3 (theoretical m/z=890.13487)
recombinant
Native m/z=890.13531, RT=9.89, Δppm=0.46 (PD)

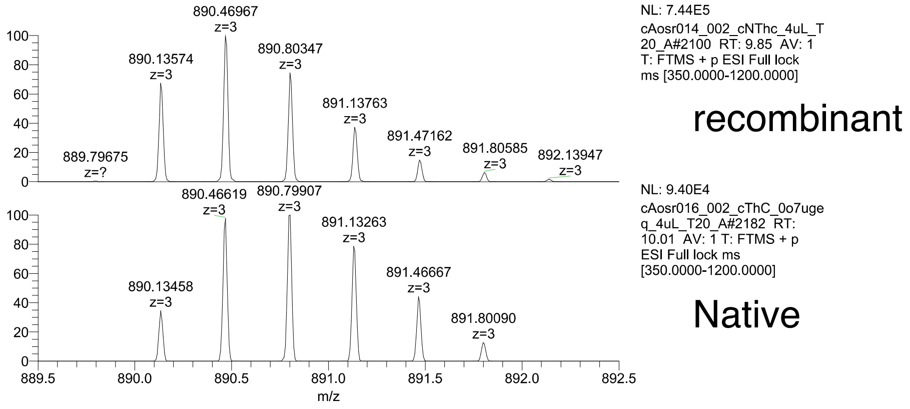

Fig. S3. B-6

ATPGILQTESSRVDNGRLIW (res 80-99)

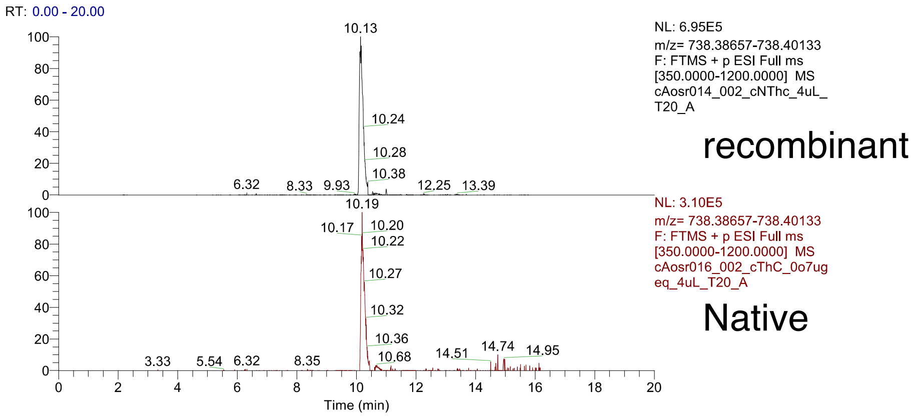

charge 3 (theoretical m/z=738.39395)

recombinant m/z=738.39441, RT=10.09, Δppm=0.57 (PD)

Native m/z=738.39429, RT=10.14, Δppm=0.41 (PD)

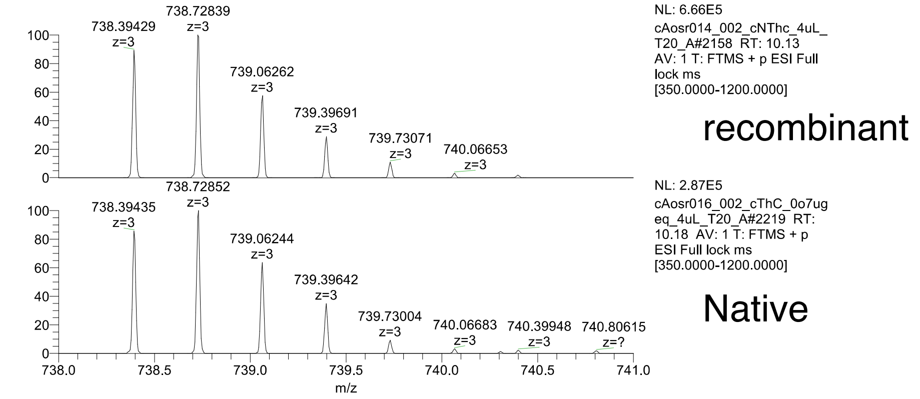

Fig. S3. B-7

### LIWQNYAFGANDGGcIVGDR (res 97-116)

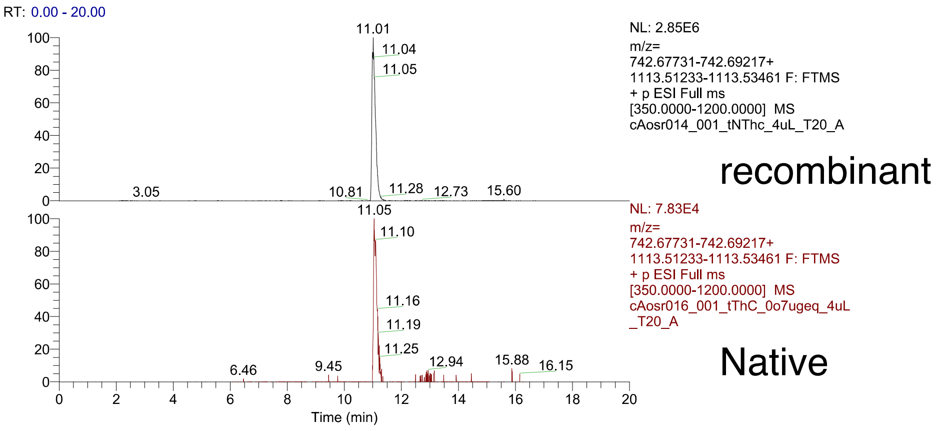

charge 3 (theoretical m/z=742.68474)

recombinant

Native m/z=742.68451, RT=11.06, Δppm=-0.36 (PD)

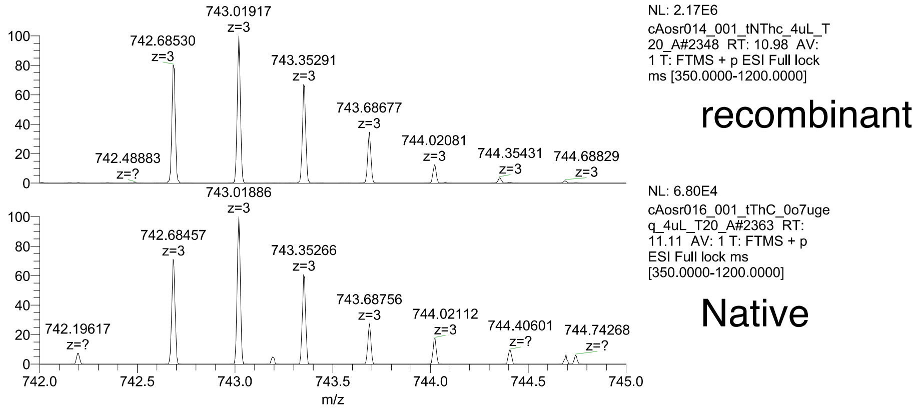

charge 2 (theoretical 1113.52347)

recombinant m/z=1113.52417, RT=10.96, Δppm=0.58 (PD)

Native

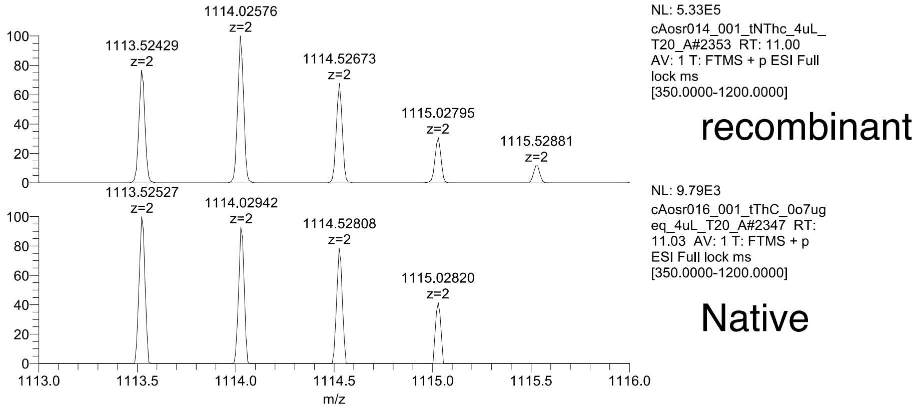

Fig. S3. B-8

GANDGGcIVGDRDFNDVF (res 105-122)

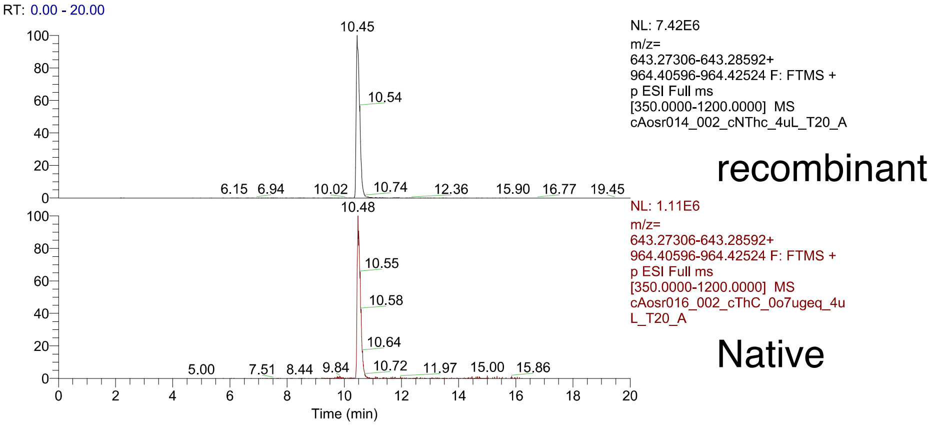

charge 2 (theoretical m/z=964.41560)
recombinant m/z=964.41559, RT=10.41, Δppm=-0.06 (PD)
Native m/z=964.41583, RT=10.43, Δppm=0.19 (PD)

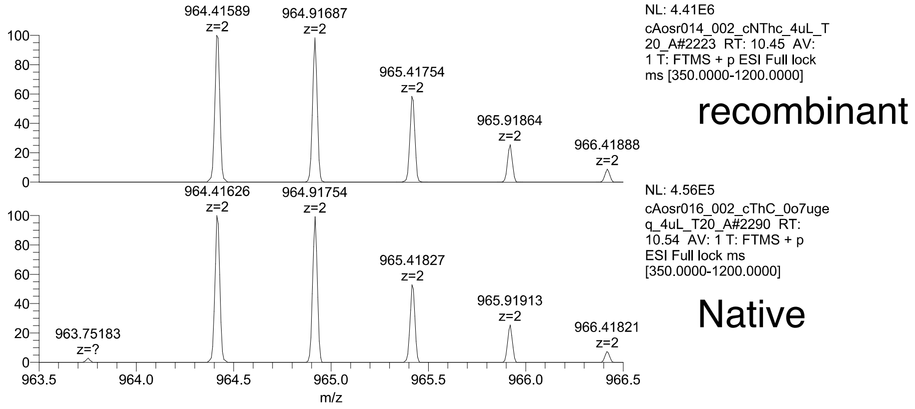

Fig. S3. B-9

### DFNDVFVLITGLVR (res 117-130)

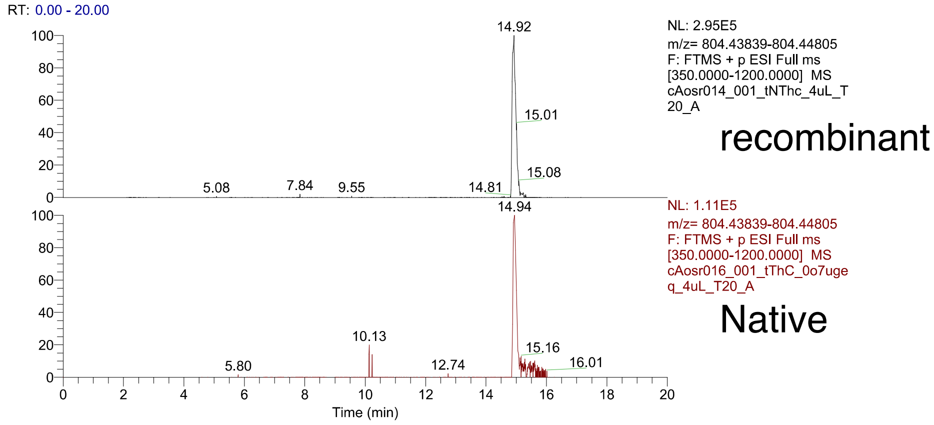

charge 2 (theoretical m/z=804.44322)
recombinant m/z=804.44360, RT=14.85, Δppm=0.42 (PD)
Native m/z=804.44287, RT=14.94, Δppm=-0.49 (PD)

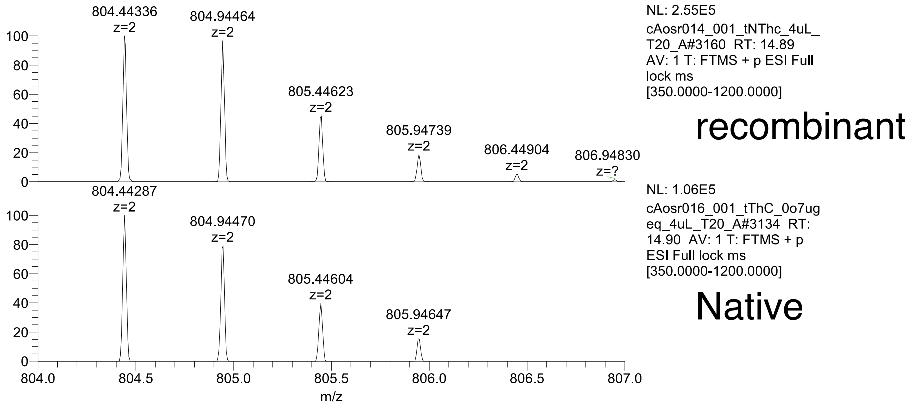

Fig. S3. B-10

### VLITGLVRG (res 123-131)

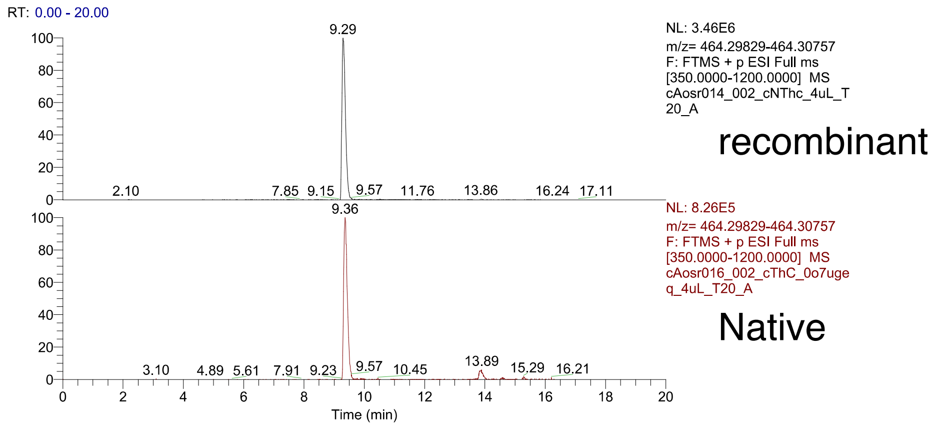

charge 2 (theoretical m/z=464.30293)
recombinant
Native m/z=464.40289, RT=9.30, Δppm=-0.14 (PD)

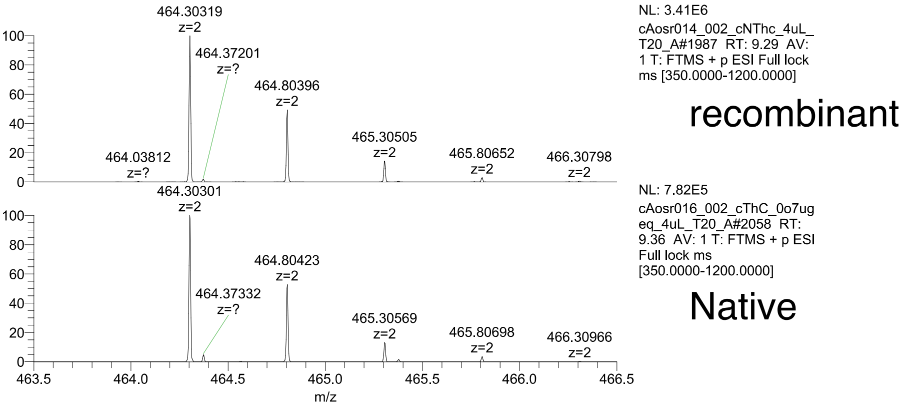

Fig. S3. C

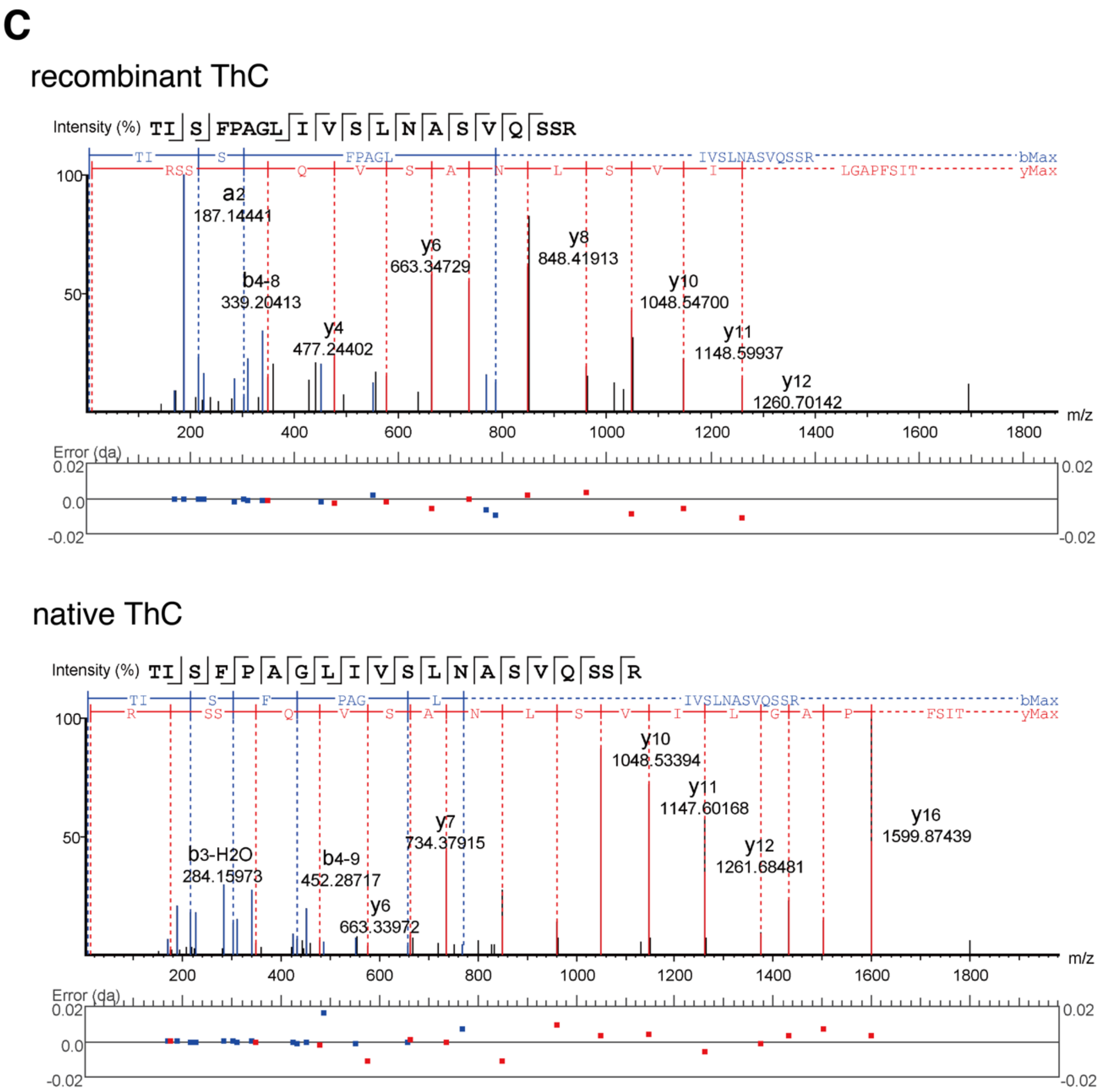

**Fig. S3. LC-MS/MS and Edman degradation mapping**

(A) Sequence coverage of nThC sequenced by LC-MS/MS. Peptide fragments, of which amino acid sequence were determined, generated by trypsin or chymotrypsin and shown red and green, respectively. Cys residue was treated with iodoacetamide to prevent from S-S bond reformation thus Cys residue was searched as Cys carbamidomethylation. (B) Mass spectrometry confirmation of the identified peptides of nThC compared with rThC. Extracted ion chromatograms (XICs) of the identified peptides of rThC and nThC were shown as black and red, respectively. MS spectra of the charged peptide ion from rThC (upper) and nThC (bottom) at the XIC peak are shown, respectively. RT, retention time; PD, the peptide identified by Proteome Discoverer (Thermo Scientific); PEAKS, the peptide identified by PEAKS studio X (Bioinformatics Solutions); ∆ppm, parts per million of the mass differences between observed and theoretical m/z values. The m/z, RT, and Δppm of the peptide identified by PD or PEAKS were indicated. Carbamidomethyl-Cys residue was shown as lower case. (C) Annotated MS/MS fragmentation spectra for a putative N-terminal 9-28 residues, TISFPAGLIVSLNASVQSSR, derived from rThC (upper panel) and nThC (lower panel). MS/MS spectra were deconvoluted into singly charged ions from the observed spectra and theoretical m/z values for fragment ions were assigned for each peak. The identified matches for the N-terminal-containing ions are shown in blue and those for the C-terminal-containing ions in red. The m/z differences between theoretical and observed values for all assigned peaks were displayed in error map and were less than 0.02 Da. The MS/MS spectra confirmed that 25$^{th}$ residue was clearly assigned as Q. The exact masses and precursor isotope distribution of MS1 spectrum for the peptides derived from nThC and rThC (1-28 residues, 25Q) was also identical (B-1), in line with above sequence analyses confirming that 25$^{th}$ residue the N terminal sequence is Q, not E.

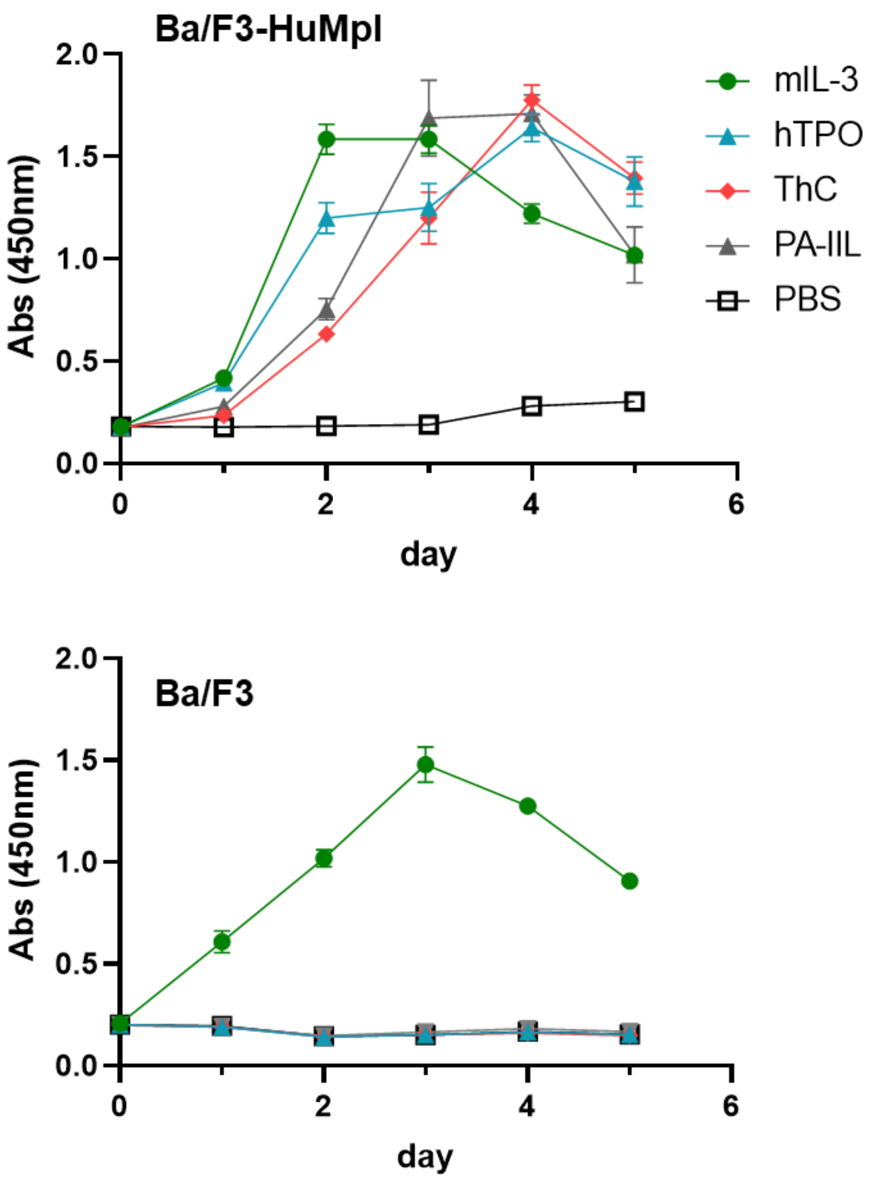

**Fig. S4. Cell proliferation assay for Ba/F3 and Ba/F3-HuMpl cells in the presence or absence of ligand**

Cell proliferation assay of Ba/F3 and Ba/F3-HuMpl cells in the absence or presence of the indicated ligand. murineIL-3 (10 ng/mL), humanTPO (10 ng/mL), ThC (0.5 μg/mL), PA-IIL (30 μg/mL).

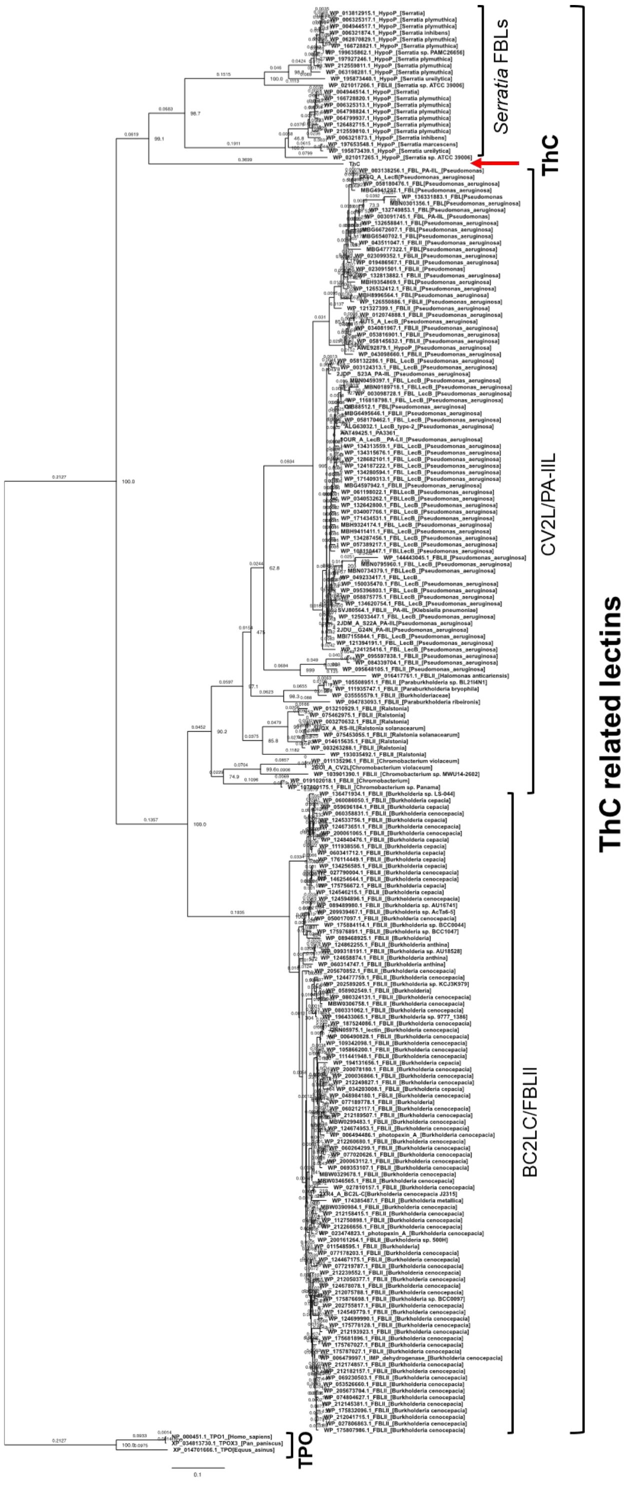

**Fig. S5. Phylogenetic tree of ThC and putative bacterial fucose binding lectins and TPO**

Bootstrap probabilities of 1,000 replications are shown at each node. A scale bar indicates the genetic distance represented by the horizontal branches.

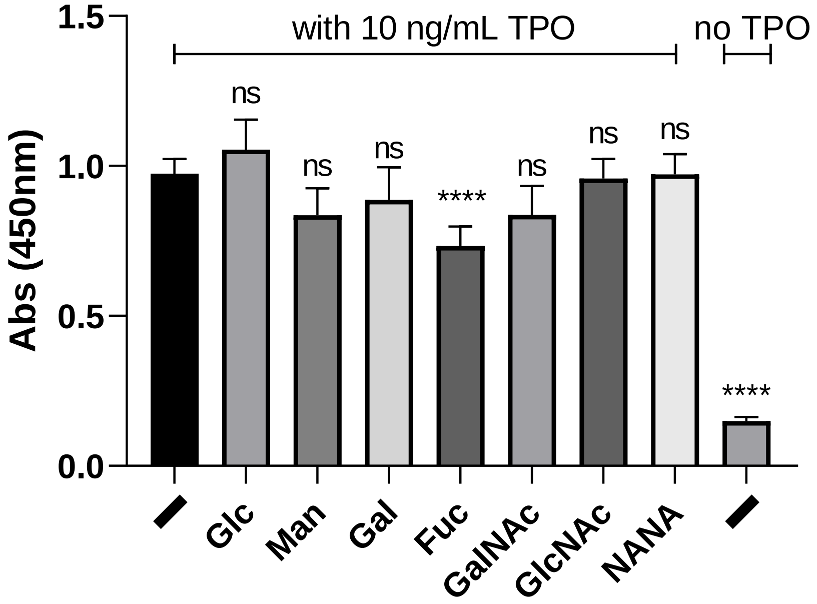

**Fig. S6. Relative cell growth by TPO in the presence of various sugars in Ba/F3-HuMpl cells**

Positive control contains TPO (10 ng/mL) and negative control without agonist. Each 10 mM of sugar was present.

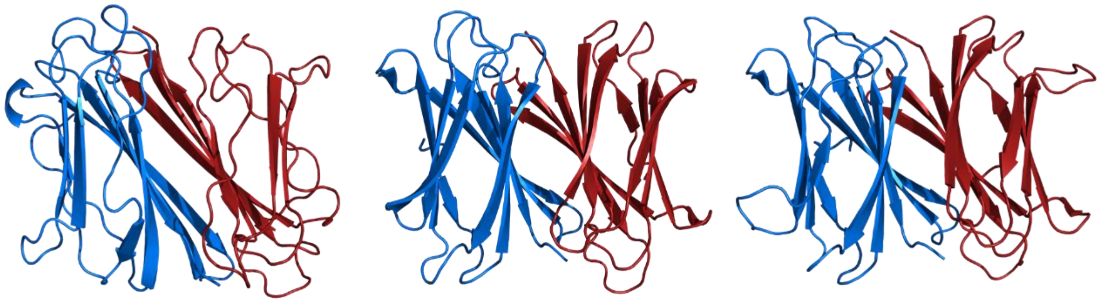

ThC dimer

PA-IIL dimer in tetramer (PDB ID 2JDP)

BC2L-C-CTD dimer (PDB ID 2XR4)

**Fig. S7. Structural comparison between ThC dimer, PA-IIL dimer (PDB ID 2JDP), and BC2L-C-CTD dimer (PDB ID 2XR4)**

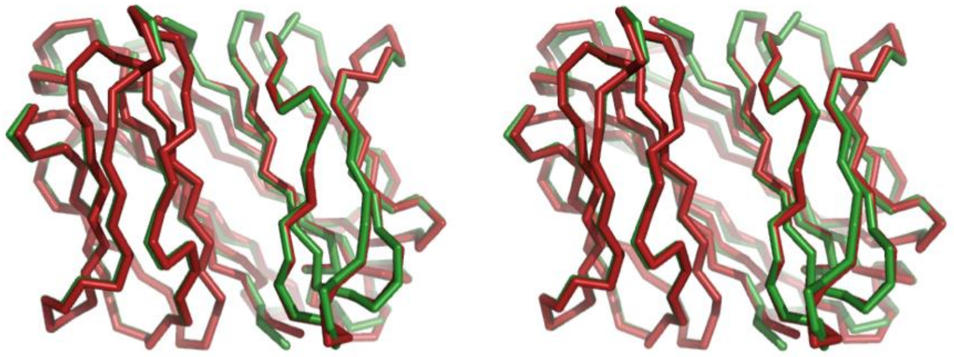

**Fig. S8. Stereo representation of superposition of nThC (green) and rThC (red)**

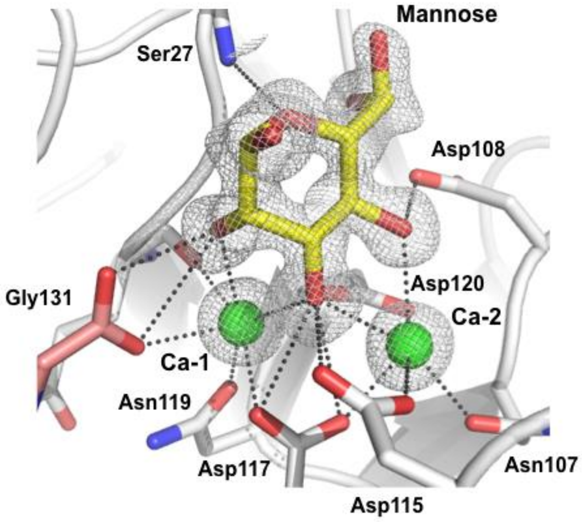

**Fig. S9. Close-up view of carbohydrate binding site in complex with rThC and D-mannose**

Fo-Fc map of the carbohydrates and $Ca^{2+}$ ions are shown. Individual protomers are shown gray and pink, respectively.

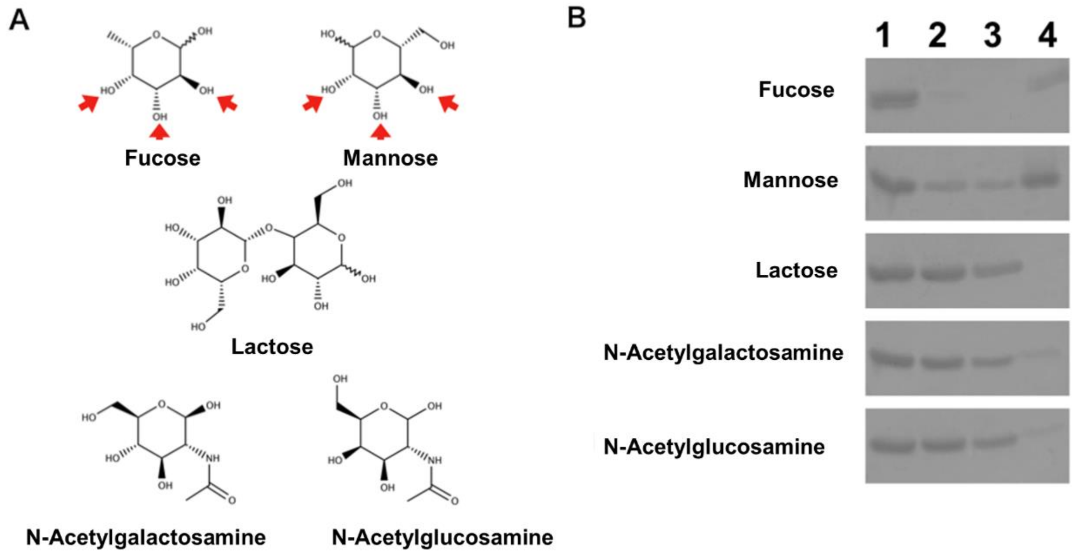

**Fig. S10. Affinity of rThC against carbohydrate-immobilized agarose gel**

Chemical structure of the carbohydrates tested (A) and their binding with rThC (B). (A) fucose, mannose, lactose, N-acetylgalactosamine, and N-acetylglucosamine. Red arrows indicate hydroxyl groups which are recognized by rThC and $Ca^{2+}$ ions. (B) Binding of rThC to the fucose-, mannose-, lactose, N-acetylglucosamine-, and N-acetylgalactosamine-immobilizing resin in the presence of 5 mM $CaCl_2$. Lane 1: rThC added to the resin, 2: flow through fraction, 3: wash fraction, 4: rThC bound on the resin.

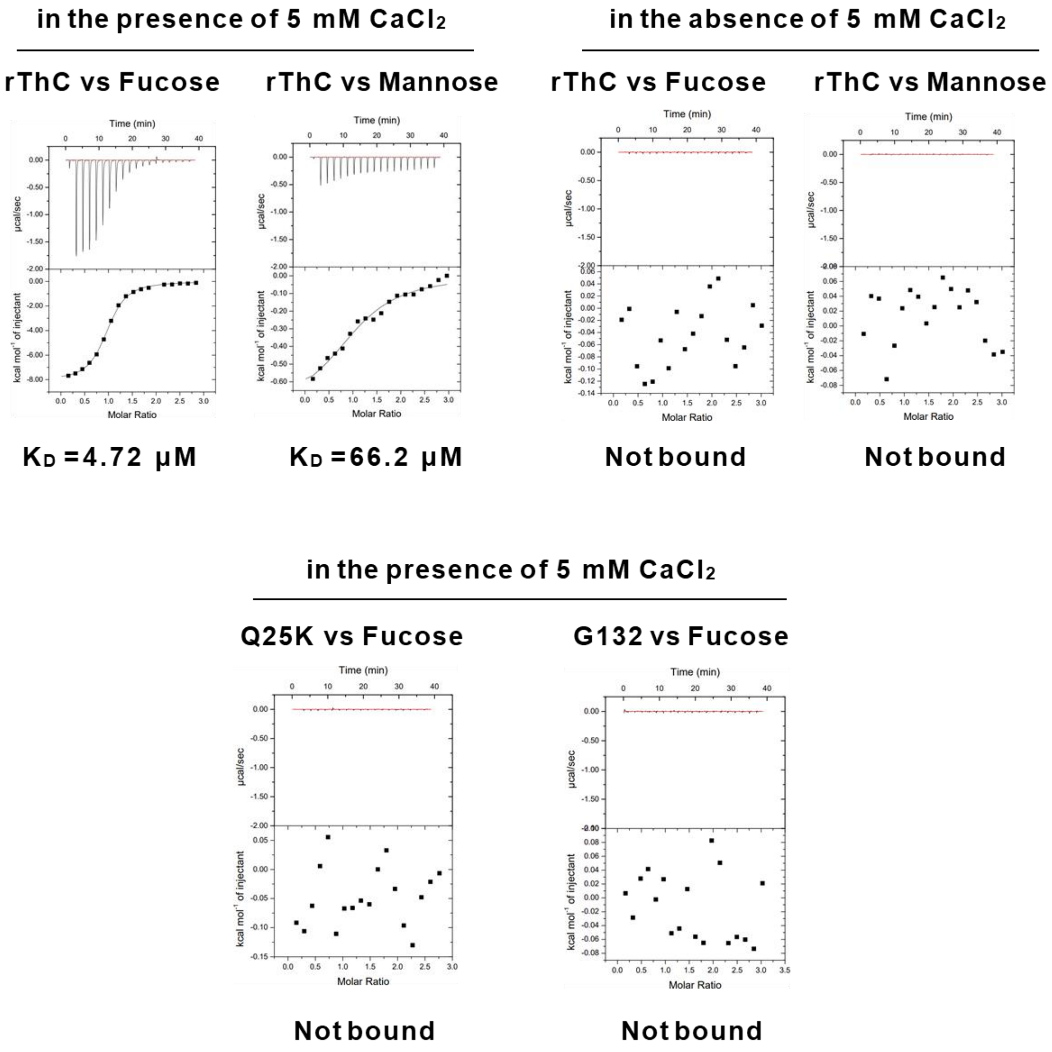

### Fig. S11. Results of ITC for rThC and mutants

rThC (top) and mutants (bottom). $K_D$ values are shown at the bottom. Thermodynamic parameters are shown in Supplementary Table S1.

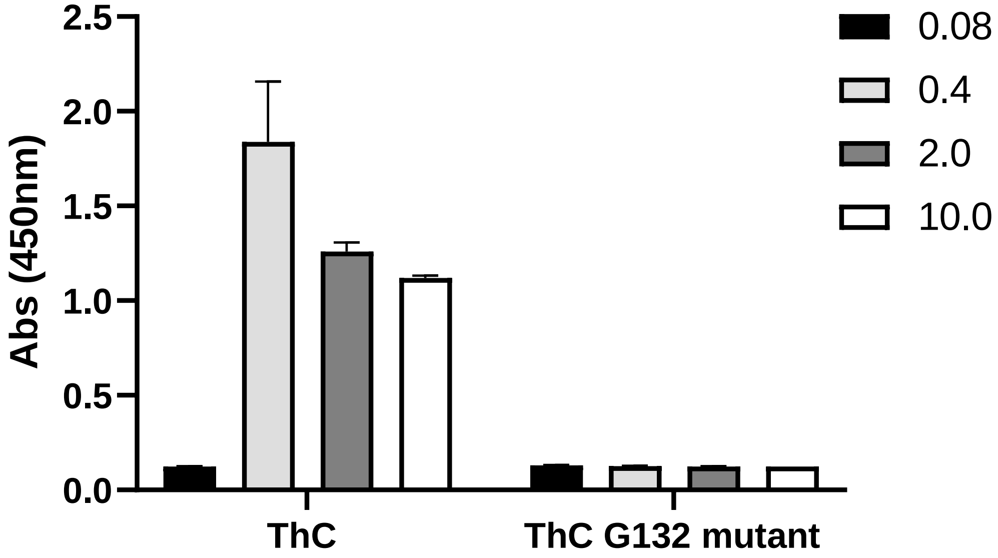

**Fig. S12. Effect of rThC G132 on cell proliferation of Ba/F3-HuMpl cells**

Cells were cultured with indicate concentration (μg/mL) ThC for 4 days.

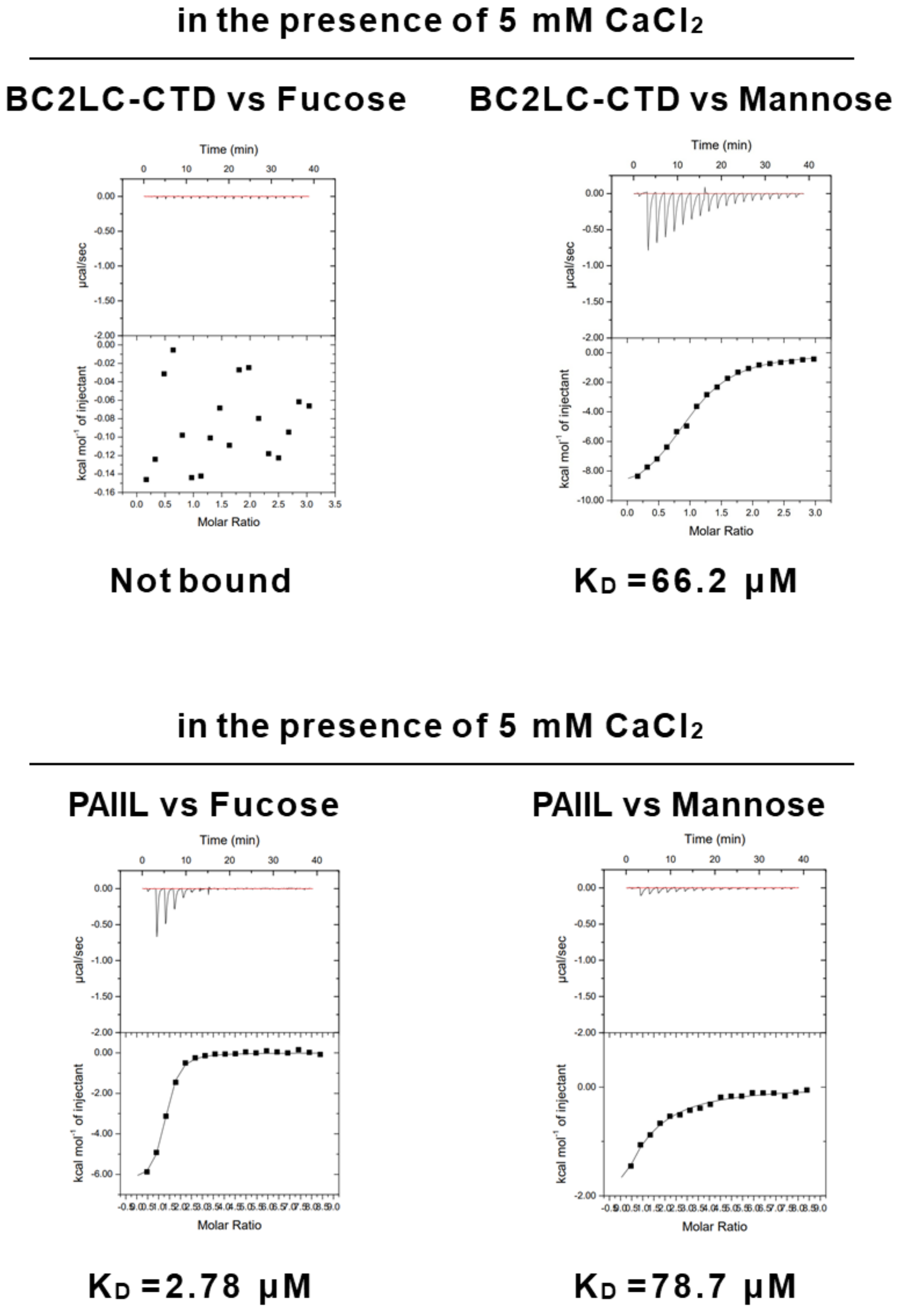

**Fig. S13. Results of ITC of BC2LC-CTD (top) and PA-IIL (bottom)**

$K_D$ values are shown at the bottom. Thermodynamic parameters are shown in Table S1.

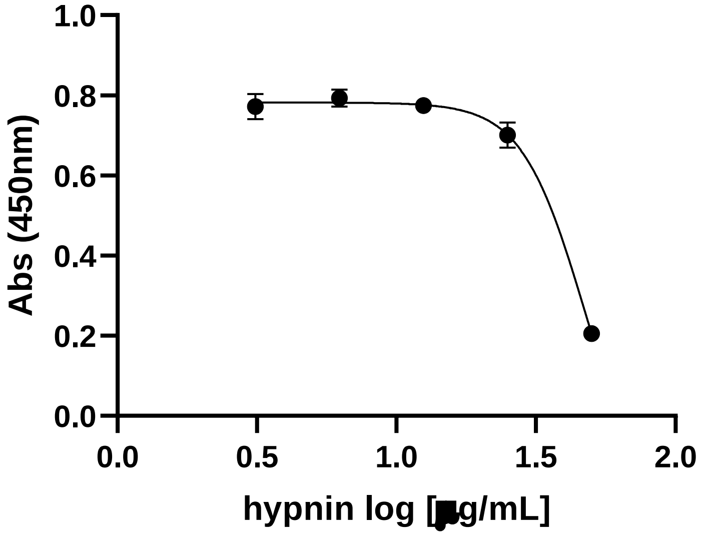

**Fig. S14. Effect of hypnin on Ba/F3 cell proliferation**

Cells were cultured with 10 ng/mL mIL-3 for 4 days. $IC_{50}$ > 47 μg/mL.

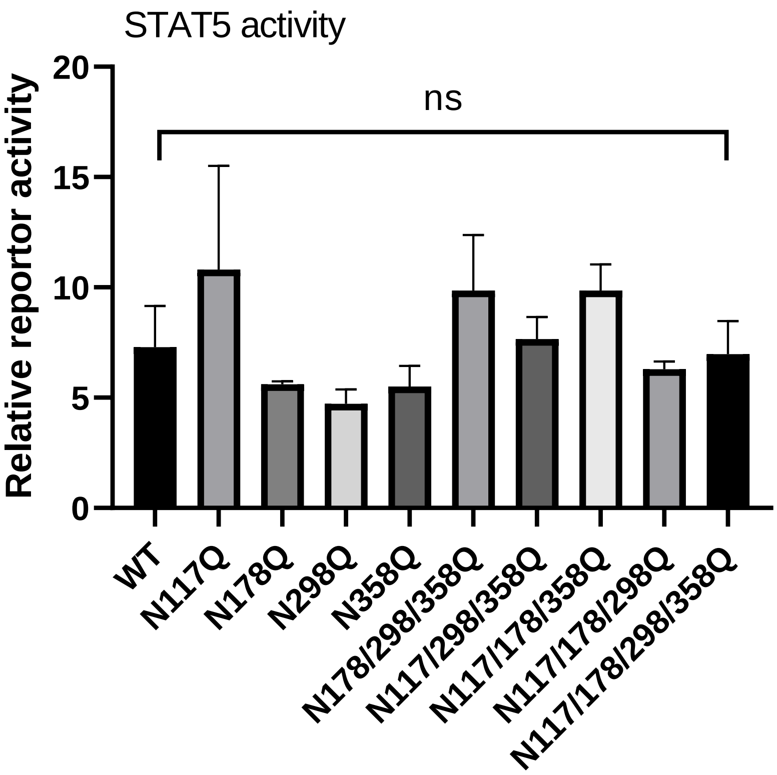

**Fig. S15. Comparable STAT5 activation by TPO with MPL bearing N/Q mutation**

STAT5 reporter activity in HEK293T cells expressing a series of N/Q mutant constructs when stimulated by TPO (10 ng/mL). ns = no significance.

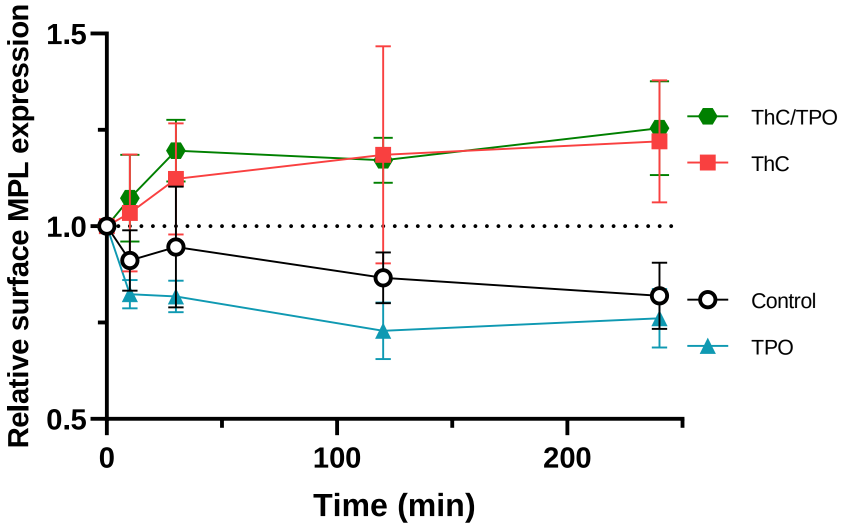

**Fig. S16. Interference of the TPO-dependent internalization of MPL from cell surface by ThC**

0, 10, 30, 120 and 240min after the addition of ThC (0.1 μg/mL), ThC/TPO (0.1 μg/mL and 0.2 ng/mL, respectively), TPO (0.2 ng/mL) and control (No agonist), levels of cell surface MPL was determined and depicted.
